# Identification of colonisation and virulence determinants of *Streptococcus pneumoniae* via experimental evolution in mouse infection models

**DOI:** 10.1101/2020.09.08.287698

**Authors:** Angharad E Green, Deborah Howarth, Chrispin Chaguza, Haley Echlin, R Frèdi Langendonk, Connor Munro, Thomas E Barton, Jay CD Hinton, Stephen D Bentley, Jason W Rosch, Daniel R Neill

## Abstract

*Streptococcus pneumoniae* is a commensal of the human nasopharynx and a major cause of respiratory and invasive disease. Pneumococcus stimulates upper respiratory tract inflammation that promotes shedding from mucosal surfaces and transmission to new hosts. Colonisation and transmission are partially antagonistic processes. Adhesion to surfaces and evasion of host responses favours the former, whilst detachment, promoted by inflammation, is necessary for the latter. We sought to determine how adaptation and evolution of pneumococcus within its nasopharyngeal niche might progress when selective pressures associated with transmission were removed. This was achieved by serial passage of pneumococci in mouse models of nasopharyngeal carriage, manually transferring bacteria between mice. To assess the role of host environmental factors on pneumococcal evolution, we also performed analogous experimental evolution in a mouse pneumonia model, passaging pneumococci through lungs. Nasopharynx-passaged pneumococci became more effective colonisers, whilst those evolved within lungs showed reduced virulence. We observed selection of mutations in genes associated with cell wall biogenesis and metabolism in both nasopharynx and lung lineages, but identified prominent examples of parallel evolution that were niche specific. We focussed on *gpsA*, a gene in which the same single nucleotide polymorphism arose in two independently evolved nasopharynx-passaged lineages. We identified a single nucleotide change conferring resistance to oxidative stress and enhanced nasopharyngeal colonisation potential. We show that *gpsA* is also a frequent target of mutation during human colonisation. These findings highlight the role played by the host environment in determining trajectories of bacterial evolution and the potential of experimental evolution in animal infection models for identification of novel pathogen virulence and colonisation factors.

## Introduction

Pneumonia and invasive disease caused by *Streptococcus pneumoniae* is a major contributor to global morbidity and mortality, but the primary lifestyle of this opportunistic pathogen is as a commensal of the nasopharynx. Compromised defence mechanisms may render a host permissive to disease, as exemplified by the rising susceptibility to pneumococcal pneumonia in the elderly^1^, whilst the suite of pneumococcal virulence factors contributes to disease development via evasion of host defences and stimulation of inflammation^2^. Several of these virulence factors are likely retained by the pneumococcus primarily as a means to ensure onwards transmission from nasopharynx, via the stimulation of the inflammation necessary to promote detachment from host surfaces.

The positive correlation between transmission and virulence is well described for a range of pathogens^3^. Experimental evolution approaches^4^ have contributed to this understanding, whereby infectious material is passaged from host to host, in an experimental system that promotes gain of infectivity via selection of individuals displaying high levels of *in vivo* growth. In studies with viral and parasite species, this approach has demonstrated that increased virulence mirrors the increase in transmissibility^5-8^.

The functional link between transmission and virulence traits has been demonstrated with pneumococcus in relation to its haemolytic toxin, pneumolysin. Pneumolysin contributes to pathology via direct lysis of host cells and stimulation of inflammation^9-11^ and pneumolysin-deficient mutants of *S. pneumoniae* demonstrate markedly attenuated virulence in animal models^9-11^. Pneumococcal lineages in which pneumolysin lacks pore-forming activity typically cause non-lethal respiratory disease in humans^12^. Zafar and colleagues have demonstrated that the inflammation stimulated by pneumolysin in the upper respiratory tract contributes to shedding of bacteria from the mucosal surfaces and thereby promotes transmission to new hosts^13^. In the absence of virus-induced inflammation, which similarly promotes shedding^14^, pneumolysin may be the primary means by which pneumococcus ensures onward transmission. However, pneumolysin is detrimental for carriage in the upper airways, as its proinflammatory properties hasten clearance of colonisation^15^. The trajectory of pneumococcal evolution has thus likely been determined by the need to balance the processes of colonisation and transmission.

We sought to determine how pneumococcus might adapt to its natural niche in the nasopharynx when selective pressures associated with transmission were removed. Transmission of *S. pneumoniae* is associated with a single-cell bottleneck^16^ and so mutations compromising transmission would not be retained in a natural setting, even if they confer advantages in colonisation. Therefore, we manually passaged pneumococci between mice in a model of asymptomatic nasopharyngeal carriage. In parallel, we also performed the same process during lung infection, to determine how environmental differences between the upper and lower airways might shape pneumococcal adaptation and evolution. Our results demonstrate rapid emergence and selection of advantageous mutations in both nasopharynx and lung passaged pneumococci. Some mutations were common to several independently-evolved lineages of pneumococcus, but these examples of parallel evolution were found in different genes in nasopharynx vs lung evolved lineages, suggesting niche-specific adaptation.

## Results

### In vivo experimental evolution of S. pneumoniae in murine nasopharynx and lungs selects for increased niche colonisation potential

To perform experimental evolution of *S. pneumoniae*, we chose the serotype 2 strain D39. Isolated nearly a century ago by Avery, this strain has been widely propagated in a laboratory environment and so may have lost many of its adaptations to the host^17^. Using an inoculum prepared from a single colony, we separately infected the nasopharynx or the lung of CD1 mice, infecting ten mice each per niche. Each of these twenty individuals was considered a founder for an independent experimental evolution lineage of pneumococcus (Figure 1). Thereafter, pneumococci were recovered from either nasopharynx or lungs. After minimal growth on agar, the entire population was used to infect a further mouse for the next round of passage. This process was repeated twenty times per lineage, with 7 days of nasopharyngeal carriage or up to 48 hours of lung infection for each round of passage. This yielded ten nasopharynx-passaged lineages and ten lung-passaged lineages. Each nasopharynx-passaged lineage had spent 140 days in total in the nasopharynx, whilst the lung-passaged lineages had spent ∼26 days in lungs.

**Figure 1.**
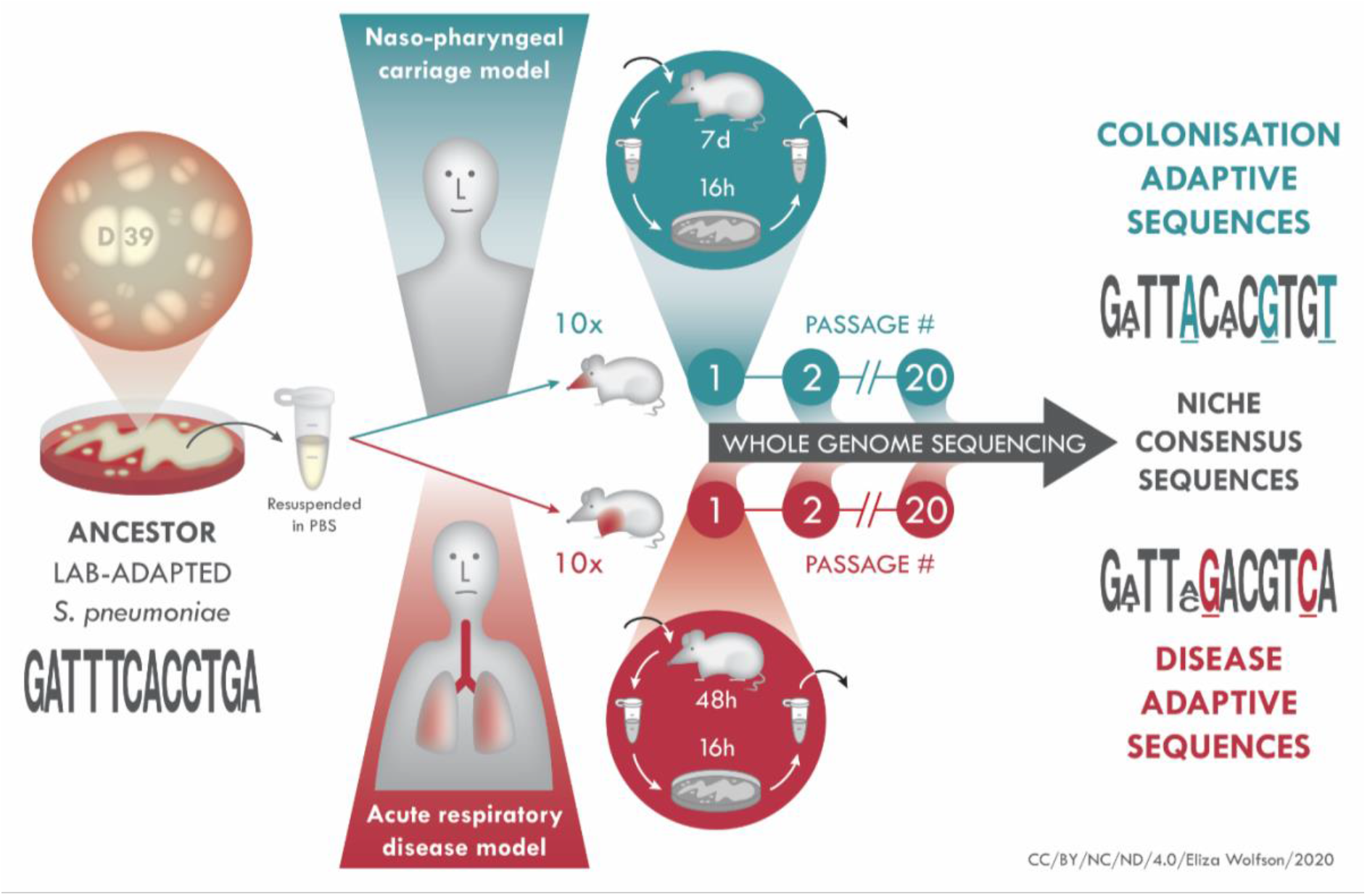
*In vivo* experimental evolution of *S. pneumoniae*. D39 (ancestor) was grown on agar, colonies resuspended in PBS and mice intranasally infected with 10 µl (carriage model, 10 mice) or 50 µl (pneumonia model, 10 mice) inoculum. The 20 infected mice were treated as founders of independent pneumococcal lineages. After each passage, which was 7 days for carriage, 48 hrs for pneumonia, pneumococci were recovered from nasopharynx, in the carriage model, or lungs, in the pneumonia model and, after minimal passage on agar, introduced into a new mouse for the next round of passage. At every passage and at the final (20x) passage, pneumococci were stored for future analysis, including whole genome sequencing for identification of adaptive mutations associated with colonisation of nasopharynx or lungs.

To determine whether *in vivo* experimental evolution had resulted in altered pneumococcal phenotypes, the twenty times nasopharynx-passaged lineages were assessed for their ability to colonise in a nasopharyngeal carriage model (Table 1, Figure 2A). All ten lineages showed increased colonisation prevalence at day 14 post-infection, relative to the ancestor and, in those mice that remained colonised, mean colonisation density was higher in all lineages except for lineage 8. (Table 1). Over a 21-day carriage experiment, nasopharyngeal colonisation density was comparable in the ancestor and a nasopharynx-passaged lineage over the first 7 days of infection, but thereafter clearance of the ancestor proceeded more rapidly (Figure 2A).

**Table 1.**
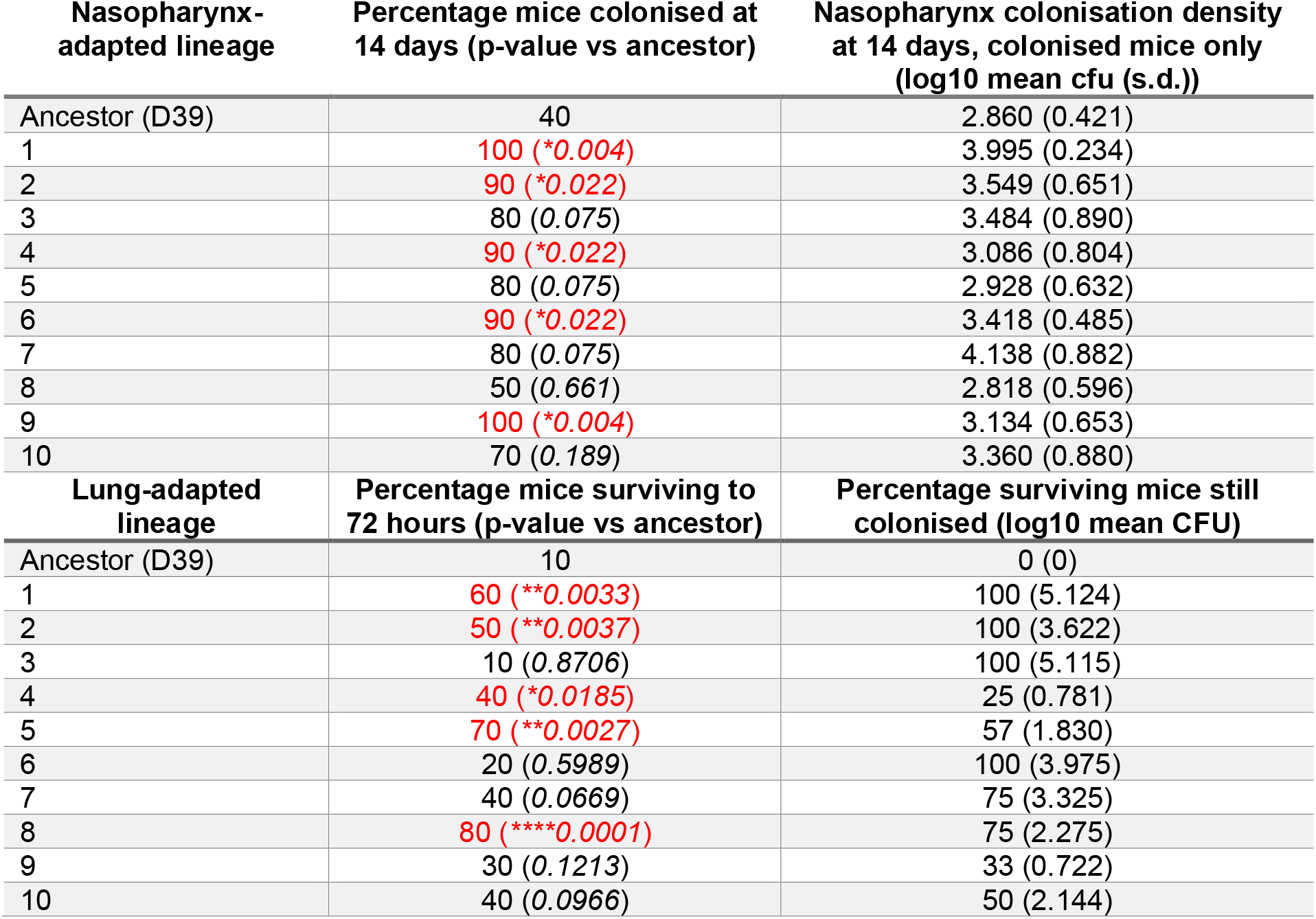
Phenotypes of nasopharynx-passaged and lung-passaged S. pneumoniae lineages. Female MF1 mice were infected with the ancestor D39 or one of the nasopharynx- or lung-passaged lineages. Nasopharynx-passaged lineages were compared in a carriage model, infecting with 1×10^5^ colony forming units (cfu) in 10 µl saline. Lung-passaged lineages were compared in a pneumonia model, infecting with 1 × 10^6^ cfu in 50 µl saline. N=10 per group for all. Colonisation was confirmed at day 14 (carriage) or day 3 (pneumonia) by plating nasopharynx or lung homogenates on blood agar. S.d. is standard deviation. P-values are from N-1 Chi squared test (carriage proportions) or Mantel-Cox log-rank test (pneumonia survival) vs the ancestor. Significant differences are shown in red.

| Nasopharynx-adapted lineage | Percentage mice colonised at 14 days (p-value vs ancestor) | Nasopharynx colonisation density at 14 days, colonised mice only (log10 mean cfu (s.d.)) |
| --- | --- | --- |
| Ancestor (D39) | 40 | 2.860 (0.421) |
| 1 | 100 (*0.004) | 3.995 (0.234) |
| 2 | 90 (*0.022) | 3.549 (0.651) |
| 3 | 80 (0.075) | 3.484 (0.890) |
| 4 | 90 (*0.022) | 3.086 (0.804) |
| 5 | 80 (0.075) | 2.928 (0.632) |
| 6 | 90 (*0.022) | 3.418 (0.485) |
| 7 | 80 (0.075) | 4.138 (0.882) |
| 8 | 50 (0.661) | 2.818 (0.596) |
| 9 | 100 (*0.004) | 3.134 (0.653) |
| 10 | 70 (0.189) | 3.360 (0.880) |
| Lung-adapted lineage | Percentage mice surviving to 72 hours (p-value vs ancestor) | Percentage surviving mice still colonised (log10 mean CFU) |
| Ancestor (D39) | 10 | 0 (0) |
| 1 | 60 (**0.0033) | 100 (5.124) |
| 2 | 50 (**0.0037) | 100 (3.622) |
| 3 | 10 (0.8706) | 100 (5.115) |
| 4 | 40 (*0.0185) | 25 (0.781) |
| 5 | 70 (**0.0027) | 57 (1.830) |
| 6 | 20 (0.5989) | 100 (3.975) |
| 7 | 40 (0.0669) | 75 (3.325) |
| 8 | 80 (****0.0001) | 75 (2.275) |
| 9 | 30 (0.1213) | 33 (0.722) |
| 10 | 40 (0.0966) | 50 (2.144) |

**Figure 2.**
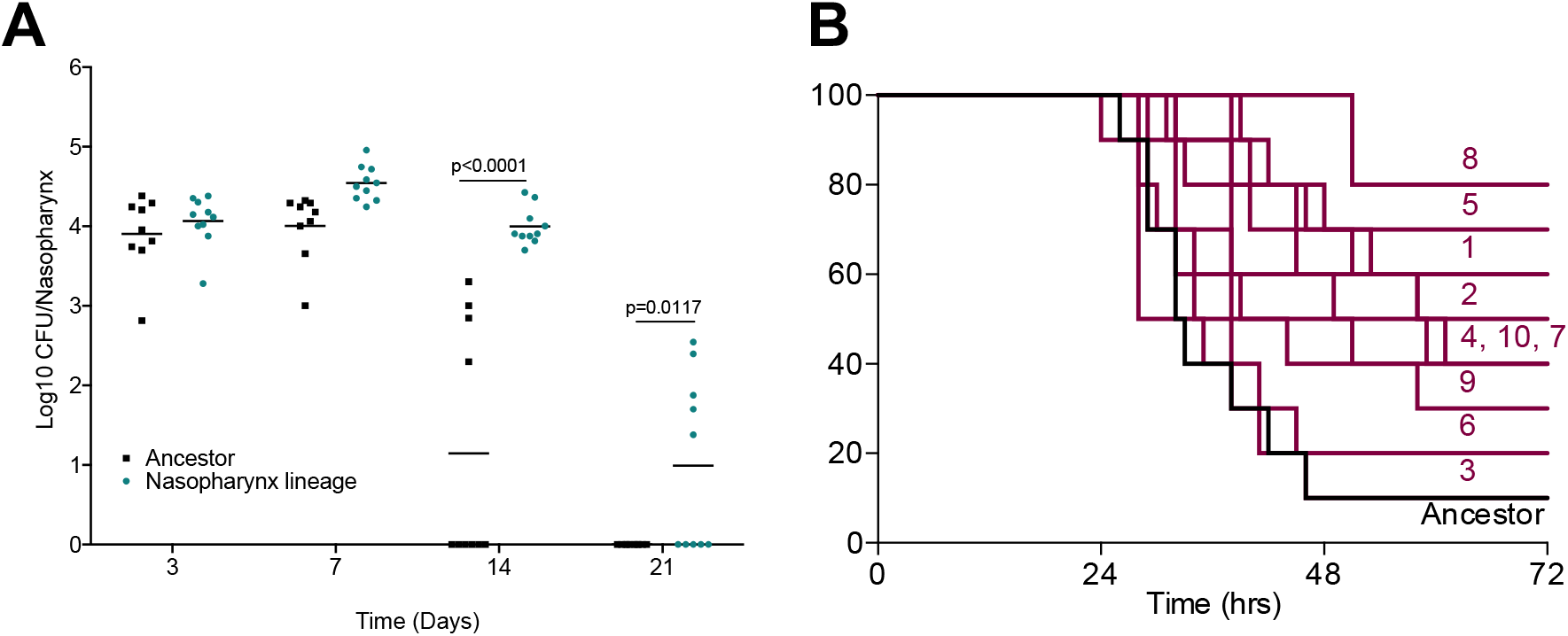
Tissue-passaged pneumococci show evidence of niche adaptation. Mice were infected with the ancestral D39 population or twenty-times passaged lineages from **(A)** nasopharyngeal carriage (lineage 1) or **(B)** pneumonia models. **(A)** Mice were infected with 1 × 10^5^ colony forming units (cfu) of the ancestor or nasopharynx lineage 1 to induce asymptomatic carriage. Bacterial numbers in nasopharynx were determined by serial dilution of tissue homogenates onto blood agar at days 3, 7, 14 and 21 post infection. P-values are from two-way ANOVA with Sidak’s multiple comparison test. **(B)** Mice were infected with 1 × 10^6^ cfu of the ancestor or one of the ten lung-passaged lineages to induce pneumonia (n=10 per group). Mice were monitored for signs of disease and culled once lethargic. Survival curves were compared by Mantel-Cox log-rank test (Table 1). Labels show lineage numbers.

Next, the twenty times passaged lung lineages were assessed for their ability to colonise murine lungs in a pneumonia model (Table 1, Figure 2B). Disease progression was slower in mice infected with lung-passaged lineages in comparison to those infected with the ancestor strain. Survival proportions were increased in mice infected with any of the ten lineages, relative to the ancestor, with significantly different survival curves for lineages 1, 2, 4, 5 and 8 (Table 1, Figure 2B). We also assessed the nasopharyngeal colonisation potential of a lung-passaged lineage and the lung colonisation potential of a nasopharynx-passaged lineage (Supplementary Figure 1). Both the nasopharynx- and lung-passaged lineages behaved comparably during lung infection, showing attenuated virulence (Supplementary Figure 1A). However, the nasopharynx-evolved lineage colonised nasopharynx at a higher density than the lung-evolved lineage over the first 14 days of infection (Supplementary Figure 1B), suggesting niche-specific fitness.

We determined growth in liquid culture, adhesion to airway epithelial cells *in vitro* and production of pneumolysin in all lineages (Supplementary Figure 2). Growth dynamics (measured as area under the logistic curve [AUC] values from 24 hours growth in broth) were comparable to that of the ancestor for all lineages, with the exception of lung-passaged lineages 7 and 8, and nasopharynx-passaged lineages 2 and 7, which displayed modestly reduced growth rates in nutrient broth (Supplementary Figure 2A and B). The levels of adhesion to A549 airway epithelial cells was highly variable in lung-passaged lineages (Supplementary Figure 2C), but two of the nasopharynx-passaged lineages showed evidence of increased adhesion potential (Supplementary Figure 2D). Levels of pneumolysin, in lysates prepared from mid-log phase pneumococci, were significantly reduced in five of ten lung-passaged lineages and nine of ten nasopharynx-passaged lineages, relative to the ancestor strain (Supplementary Figure 2E and F). The level of epithelial cell adhesion was negatively correlated with the amount of pneumolysin produced in lung passaged lineages (R^2^ = 0.70) (Supplementary Figure 2G).

### Genomic characterisation of niche-passaged pneumococci

To uncover the genetic basis of the observed adaptations, we genome sequenced 100 pneumococcal populations, which included all twenty times passaged lineages and passage numbers 1, 5, 10 and 15 of each lineage. By comparison with the genome sequence of the ancestor strain, we identified variants that had arisen during experimental evolution. We first performed *de novo* genome assembly for our ancestor D39, using a combination of long- and short-read sequencing. This yielded a single contig genome of 2,046,551 base pairs, a size similar to that determined in a recently published assembly for this strain^18^. Genome annotation with Prokka^19^ identified 1,998 protein coding sequences, with 664 of these being hypothetical proteins. Deep sequencing of the ancestor population (300x coverage) was performed to enable us to detect low frequency variants in the inoculum used to establish the 20 experimental evolution lineages. Using Breseq^20^, we mapped the Illumina reads from the ancestor inoculum against the consensus genome sequence. After filtering out artefactual or unreliable mutations and those at <2% frequency (the suggested frequency-cut off limit for polymorphism mode in Breseq^20^), we observed 43 unique variants, at frequencies of between 2.2 and 20.7% in the total ancestor population (Supplementary Dataset 1). Most of these were subsequently lost during *in vivo* experimental evolution. Those that were retained (in genes *dnaK* and *P0_01974*) are highlighted below.

Variant calling of Illumina reads from *in vivo* passaged pneumococci, with an average of 147x coverage (range 105x-190x), detected a total of 735 variants across all the nasopharynx lineages (passages 1, 5, 10, 15 and 20), with 206 variants identified in the final populations that had been passaged twenty times through nasopharynx (Supplementary Dataset 1). Within lung-passaged lineages we observed 509 variants, with 128 found across the ten populations at passage twenty (Supplementary Dataset 1). The majority of variants detected were SNPs, although deletions of between 1 and 53 base pairs were identified, as well as several single nucleotide insertions. Indels were predominantly located in intergenic regions. Cluster of orthologous gene (COG) analysis^21^ of variant data demonstrated that genes with roles in amino acid and nucleotide metabolism accumulated mutations in lineages from both niches. In lung-passaged lineages, 16% of variants were found in genes associated with carbohydrate transport and metabolism, whilst 6% of variants in nasopharynx-passaged lineages were in genes associated with energy production and conversion (Supplementary Figure 3).

**Figure 3.**
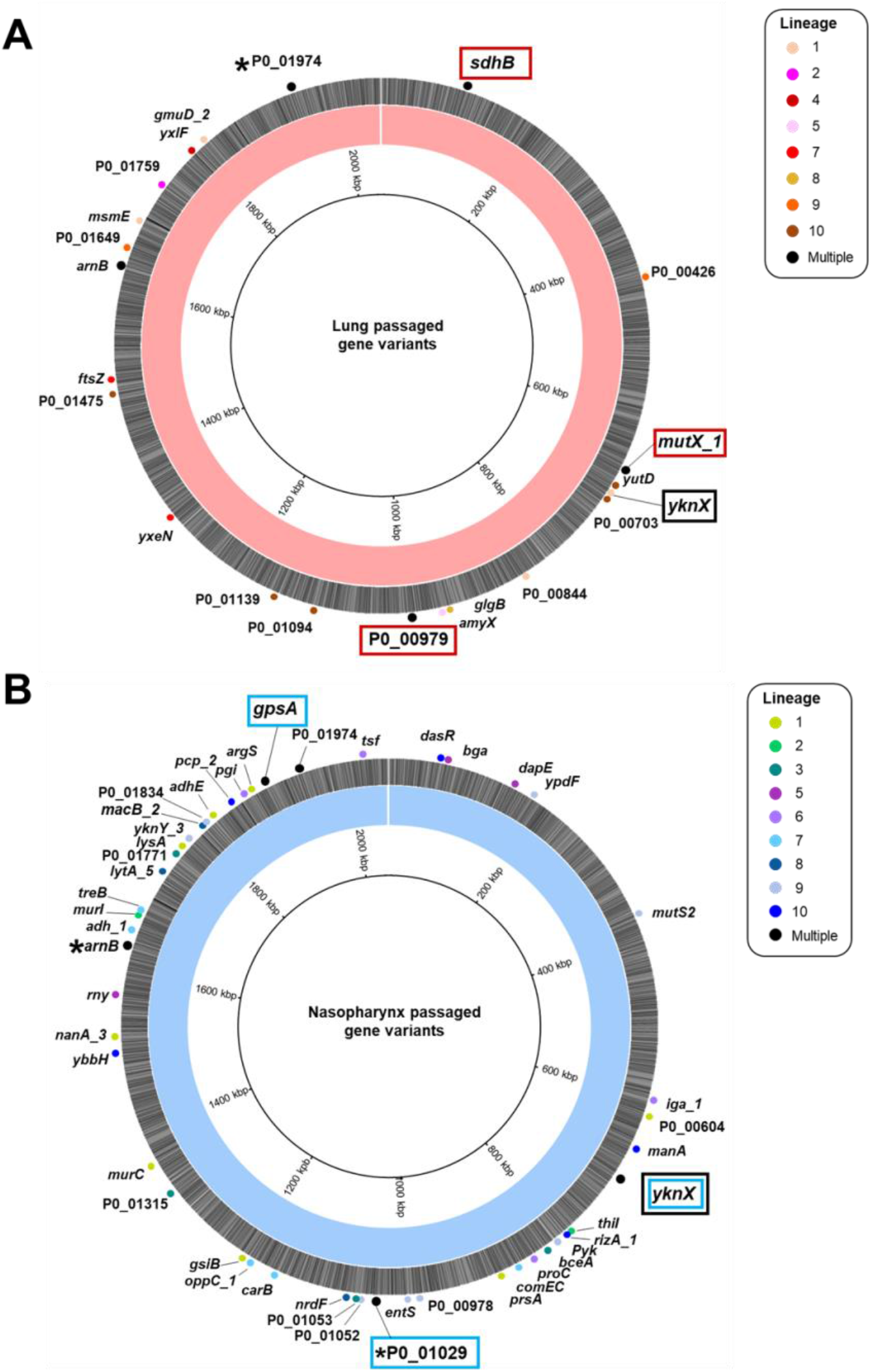
Fixed non-synonymous mutations identified in twenty-times passaged *S. pneumoniae* lineages evolved in mouse lungs or nasopharynx. Genome maps of the D39 ancestor strain assembly, highlighting the non-synonymous single nucleotide polymorphisms identified in twenty-times passaged pneumococci from the lung **(A)** or nasopharynx **(B)**. Labels show the name and position of the gene in which the SNP occurs, with the circles coloured according to the lineage where the variant was identified. Black indicates SNPs identified in multiple lineages. Potential examples of parallel evolution are highlighted in red boxes for the lung and blue boxes for the nasopharynx evolved lineages. The same SNP in the *yknX* gene was identified in two nasopharynx lineages, with a separate SNP identified in one lung lineage. This gene is highlighted with a black box. The mutations for all genes are defined in Supplementary Dataset 1 and *’s indicate those genes where different mutations were observed in separate lineages. Variants in *arnB* and *P0_ 01974* were also identified in the ancestor and control passage.

To identify mutations most likely to drive adaptive phenotypic change, we focused on mutations that became fixed in the population during experimental evolution. Considering only the final populations in each lineage (that is, those had been passaged twenty times through a niche), we identified 47 different non-synonymous SNPs within nasopharynx-passaged pneumococci (Figure 3A), and 31 in lung-passaged pneumococci (Figure 3B), that became fixed (reached 100% frequency) in at least one lineage. These included several examples of potential parallel or convergent evolution, whereby the same mutation, or closely located mutations, arose and reached fixation in two or more independently evolved lineages (Table 2). Two of these were observed only in nasopharynx-passaged lineages, three were observed only in lung-passaged lineages and three were found in lineages from both nasopharynx and lungs. Despite the notable changes in pneumolysin production observed, none of the lung or carriage lineages had any variants in the *ply* gene or flanking regulatory regions.

**Table 2.**
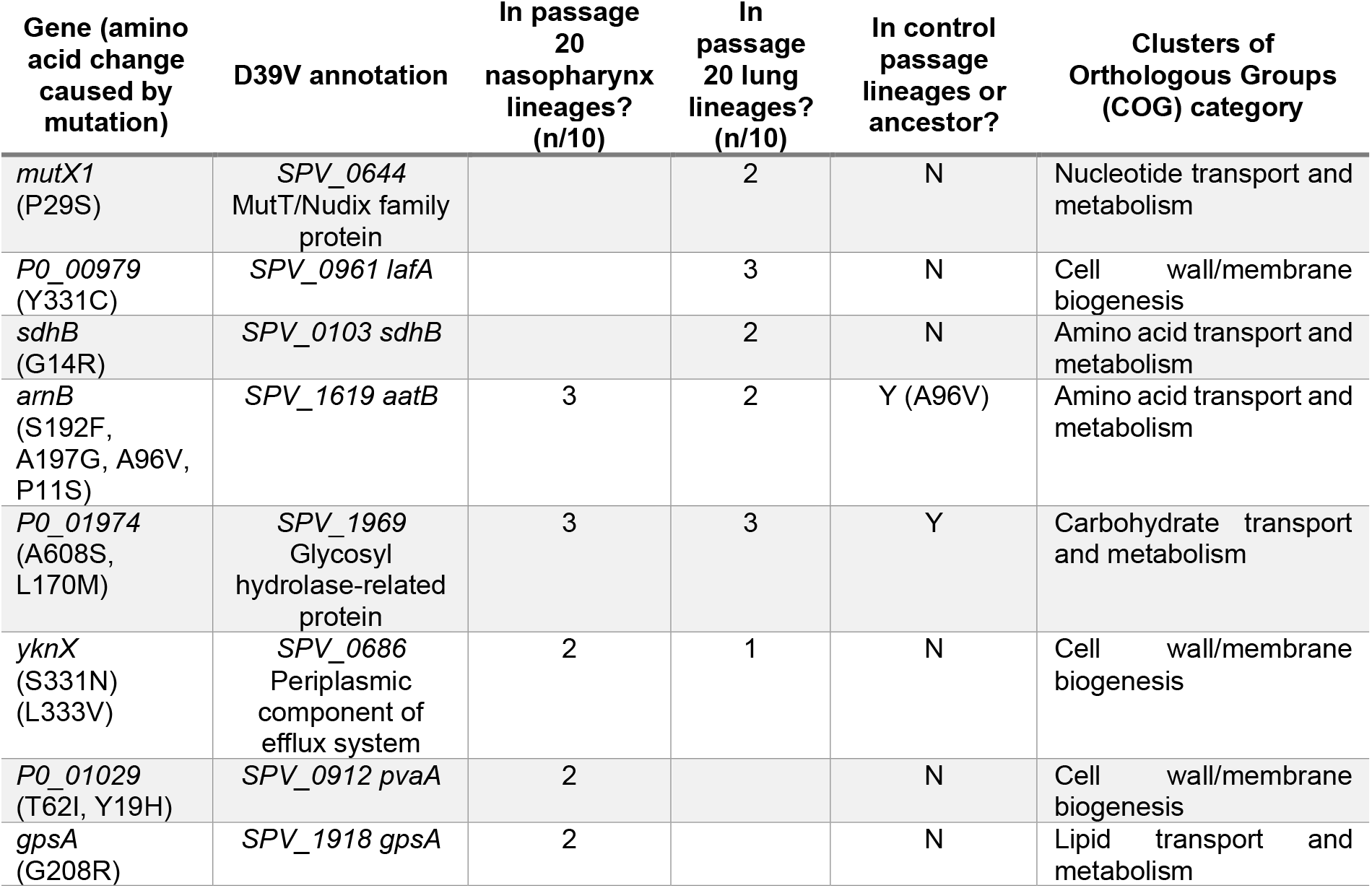
Parallel or convergent evolution in niche-passaged pneumococcal lineages. Table shows all genes in which non-synonymous single nucleotide polymorphisms were identified in more than one independently evolved lineage. Columns, from the left, show; 1. the gene affected and the resulting amino acid change(s), 2. the corresponding gene in the D39 genome recently published by Slager and colleagues^18^, 3. the number of twenty-times nasopharynx passaged lineages in which the variant is found at 100% frequency, 4. the number of twenty-times lung passaged lineages in which the variants is found at 100% frequency, 5. whether the same variant was identified in either the ancestor population or the laboratory passaged (control) lineages, and 6. The COG classification^21^ of the affected gene.

| Gene (amino acid change caused by mutation) | D39V annotation | In passage 20 nasopharynx lineages? (n/10) | In passage 20 lung lineages? (n/10) | In control passage lineages or ancestor? | Clusters of Orthologous Groups (COG) category |
| --- | --- | --- | --- | --- | --- |
| *mutX1* (P29S) | SPV_0644 MutT/Nudix family protein |  | 2 | N | Nucleotide transport and metabolism |
| *P0_00979* (Y331C) | SPV_0961 *lafA* |  | 3 | N | Cell wall/membrane biogenesis |
| *sdhB* (G14R) | SPV_0103 *sdhB* |  | 2 | N | Amino acid transport and metabolism |
| *arnB* (S192F, A197G, A96V, P11S) | SPV_1619 *aatB* | 3 | 2 | Y (A96V) | Amino acid transport and metabolism |
| *P0_01974* (A608S, L170M) | SPV_1969 Glycosyl hydrolase-related protein | 3 | 3 | Y | Carbohydrate transport and metabolism |
| *yknX* (S331N) (L333V) | SPV_0686 Periplasmic component of efflux system | 2 | 1 | N | Cell wall/membrane biogenesis |
| *P0_01029* (T62I, Y19H) | SPV_0912 *pvaA* | 2 |  | N | Cell wall/membrane biogenesis |
| *gpsA* (G208R) | SPV_1918 *gpsA* | 2 |  | N | Lipid transport and metabolism |

We also performed a control passage process, whereby the ancestor was serially passaged 20 times on blood agar, to control for any selective pressures associated with sequential growth on agar plates during evolution of the niche-evolved lineages. Variants identified in control passage lineages largely differed from those identified in the nasopharynx and lung lineages (Supplementary Dataset 1). Of the SNPs in Table 2, the *P0_01974* mutation was identified in both the ancestor population and 6/8 control passage lineages, and one of the four identified *arnB* mutations (A96V) was found in 7/8 control passage lineages. All SNPs in Table 2 that were present in lung or nasopharynx lineages but were not found in the ancestor or control passage were subsequently confirmed by Sanger sequencing.

### Evidence of parallel evolution in niche-passaged pneumococci

Three lineages of lung-passaged pneumococci acquired the same SNP in *P0_00979*, encoding the glycosyltransferase LafA, involved in synthesis of lipid anchors for cell wall teichoic acid^22^. This SNP arose early in lung lineages 4, 5 and 10 and quickly reached fixation within the population, as can be seen in Muller plots depicting genotype frequencies and ancestries (Figure 4A, B). This SNP was not detected in the ancestor or any of the nasopharynx or control passage lineages, suggesting it could confer an advantage within the lung environment.

**Figure 4.**
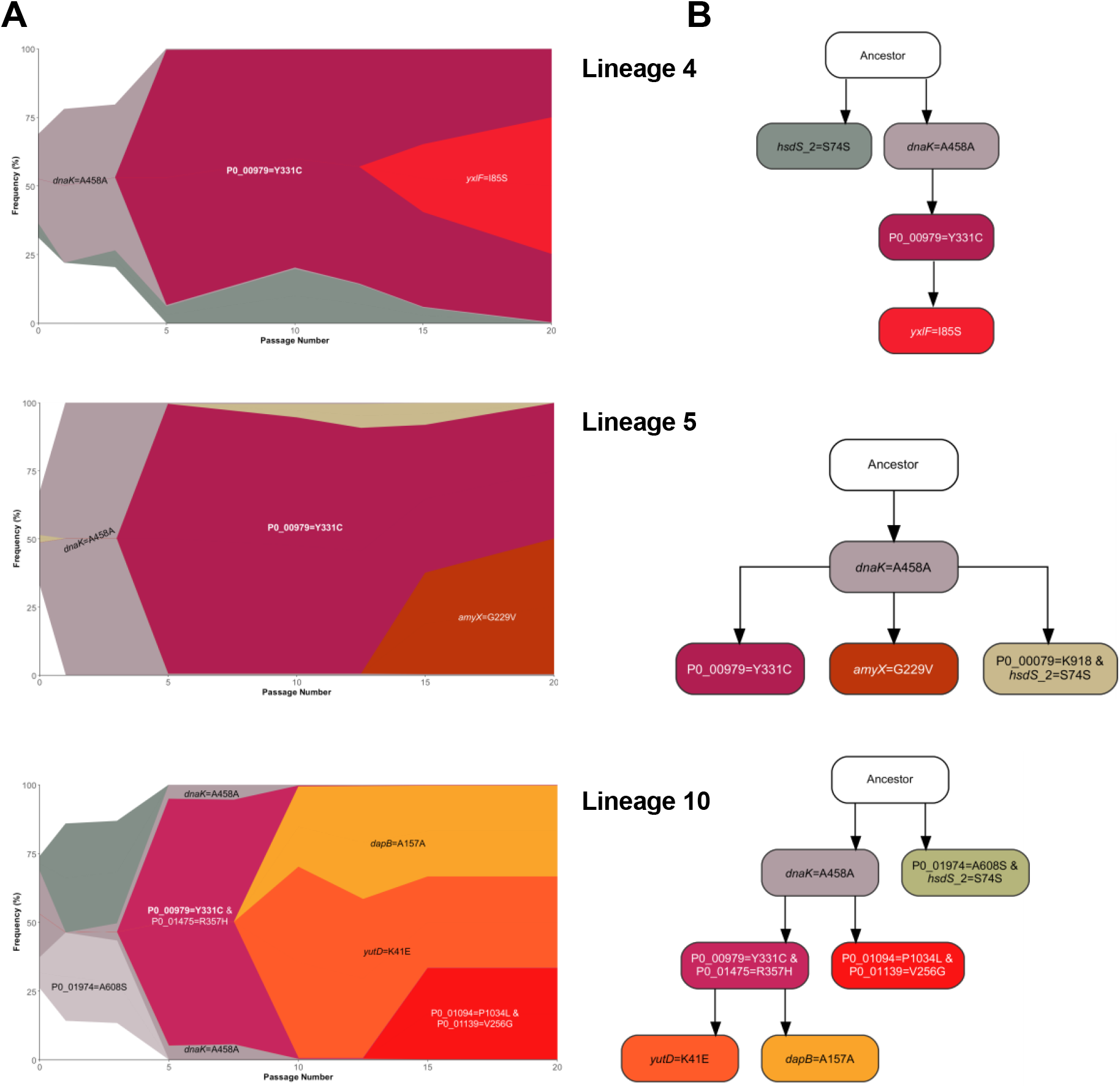
Success of *lafA* polymorphisms in lung-passaged pneumococcal lineages. **(A)** Muller plots showing genotype ancestry and frequency for lineages 4, 5 and 10 of lung-passaged pneumococci. The genotype associated with the *P0_00979* (*lafA*) SNP is shown in maroon. For lineages 4 and 5, this genotype is uniquely associated with this SNP, whilst in lineage 10 the genotype is also defined by a non-synonymous mutation in *P0_01475*. Additional colours are arbitrary but other successful genotypes containing non-synonymous SNPs are labelled. The ancestral genotype is in white. **(B)** Ancestry diagrams inferring relationships between genotypes within the population. Where multiple variants define a genotype, order of acquisition is uncertain.

For nasopharynx-passaged lineages, two examples of potential parallel evolution were found in the mutations in *P0_01029* found in lineages 1 and 7, and the co-occurring *gpsA* and *yknX* SNPs in lineages 3 and 4. Lineages 1 and 7 each acquired a different mutation in *P0_01029* but both resulted in an amino acid change, at positions 62 and 19, respectively. This gene encodes the surface-expressed pneumococcal vaccine antigen A (*pvaA*), identified in an immunising screen of pneumococcal proteins as affording protection against disseminated infection^23^. Hava and Camilli highlighted this gene as being essential for lung infection^24^. The mutations in *pvaA* arose early in both lineages and quickly reached fixation (Figure 5A and B). Carriage lineages 3 and 4 both acquired the same non-synonymous mutations in *gpsA*, encoding a glycerol-3-phosphate dehydrogenase, and *yknX*, encoding a component of an uncharacterised efflux pump. These SNPs became fixed in the populations (Figure 5C). Muller plots, genotype ancestries and genotype frequency plots for all lung and nasopharynx passaged lineages can be found in Supplementary Figures 4-13.

**Figure 5.**
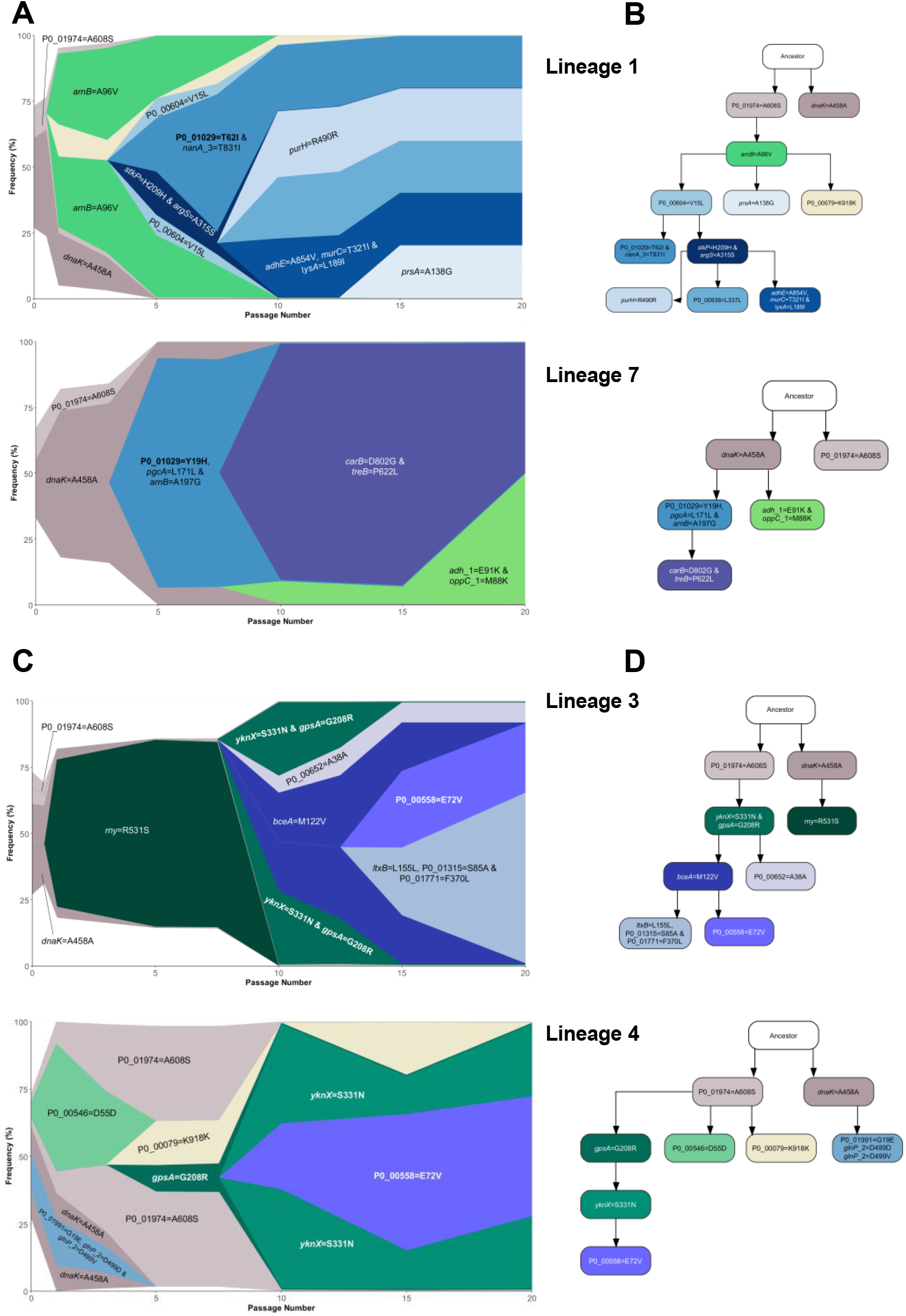
Parallel evolution in nasopharynx-passaged pneumococci. **(A)** Muller plots showing the emergence and success of *P0_01029* (*pvaA*) variants in nasopharynx-passaged lineages 1 and 7. The *pvaA-*associated genotype is shown in teal. Additional colours are arbitrary but other successful genotypes or those originating from the ancestor population are labelled. **(B)** Genotype ancestry for lineages 1 and 7. **(C)** Muller plots showing the emergence and success of *gpsA* and *yknX* variants in nasopharynx-passaged lineages 3 and 4. The *gpsA* and *yknX* genotypes are shown in shades of green. The two variants are associated with a single genotype in lineage 3. A shared non-synonymous variant in *P0_00558* is shown in purple. Additional colours are arbitrary. **(D)** Genotype ancestry for lineages 3 and 4.

Several common trends were observed amongst lineages. Between 2 and 22 distinct variants were detected per lineage, per passage, in pneumococci from lungs (Figure 6A). Between 4 and 29 distinct variants were detected per lineage, per passage in nasopharynx-passaged pneumococci (Figure 6B). Both lung and nasopharynx passaged lineages had increasing variant numbers in later passages (*p<0*.*0001* at passage 15 vs passage 1, *p=0*.*0057* at passage 20 vs passage 1 for lung lineages, *p=0*.*0113* at passage 15 vs passage 1, *p<0*.*0001* at passage 20 vs passage 1 for nasopharynx lineages) (Figure 6C). The mean number of variants detected was lower at each passage in lung lineages vs nasopharynx, perhaps reflecting their reduced time spent *in vivo*. Most variants identified were detected only once, that is, in a single lineage at a single passage. However, there was evidence of selection for some SNPs, with ten appearing ≥10 times in nasopharynx lineages and eleven appearing ≥10 times in lung lineages (Figure 6D). Of the total unique variants identified, discounting those present in the ancestor population, 245 were found only in nasopharynx lineages and 118 were found only in lung lineages (Figure 6E). Within the 22 variants shared between nasopharynx and lung populations, 13 were also found in control passage lineages.

**Figure 6.**
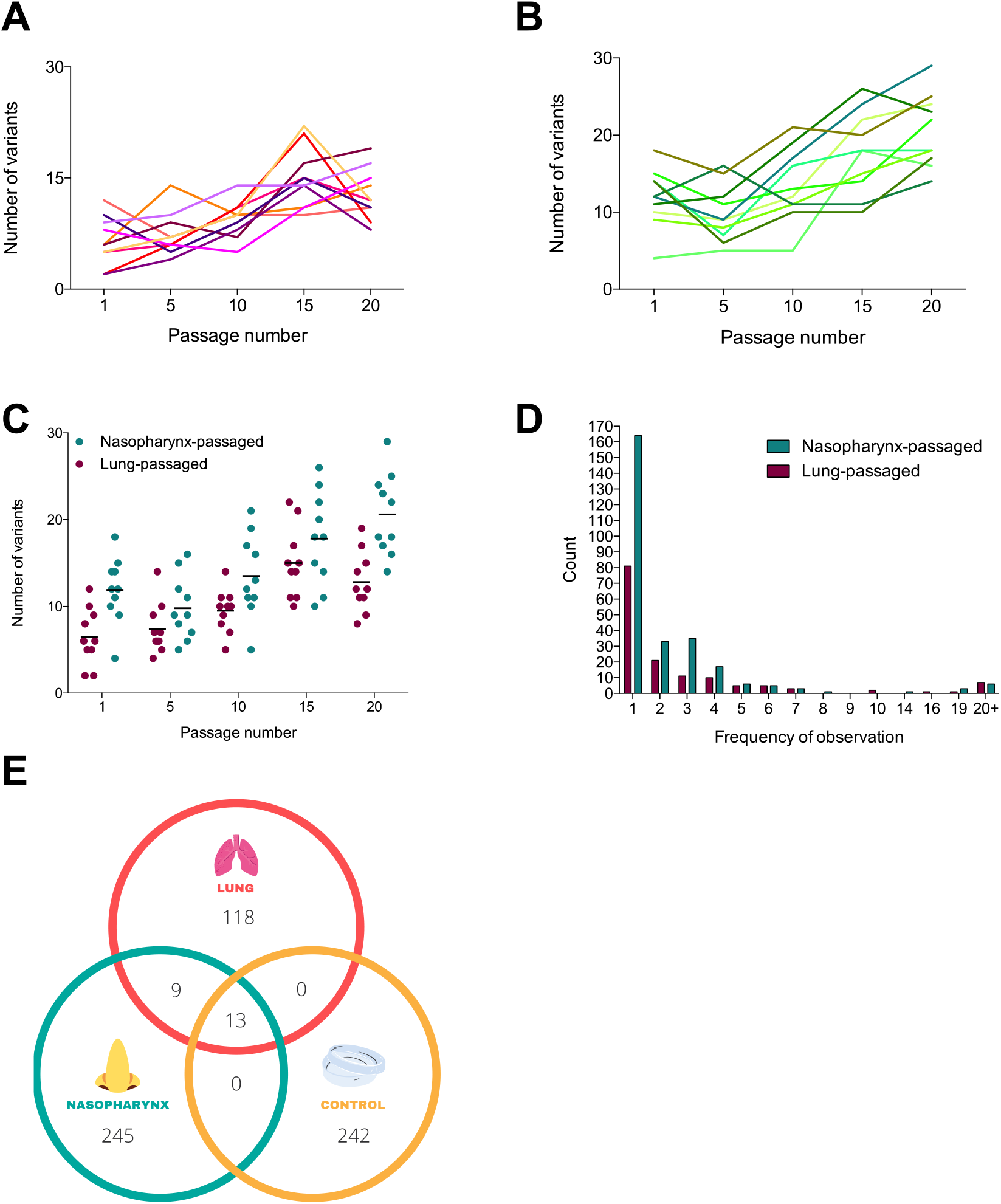
Variant dynamics in niche-passaged pneumococci. **(A)** Number of variants identified per passage in ten *S. pneumoniae* lineages during passage through murine lungs in a pneumonia model. Each line represents an independently evolving lineage (n=1 per lineage, per timepoint). **(B)** Variants per passage for ten independently evolving nasopharynx-passaged lineages in a murine carriage model (n=1 per lineage, per timepoint). **(C)** Comparison of variant numbers per passage in nasopharynx and lung passaged pneumococci. Each dot represents data from a single lineage. Data are a composite of those in **(A)** and **(B)** and are included for clarity. **(D)** Number of times each unique variant was detected. Data are from a composite of all passages of all lineages. For example, if the same variant is detected at passage 1 and 5 in lineage 2 and at passage 20 in lineage 3, but is not found in other lineages, it would be recorded as a count of 3. **(E)** Venn diagram showing unique variants in nasopharynx-, lung-, or lab (control) passaged lineages. Here, each variant is counted only once, regardless of the number of times it was detected.

### Function of gpsA, a gene associated with adaptation to the nasopharynx

We observed a guanine to adenine substitution in *gpsA*, resulting in a glycine to arginine transition at amino acid 208, in nasopharynx-passaged lineages 3 and 4. We focussed on this gene, because in a recent longitudinal study, assessing microevolution of pneumococcus over 12 months in colonised infants, SNPs in *gpsA* were observed at a frequency of 35.36 per kilobase pair, placing it 23^rd^ out of 592 genes in which SNPs were detected, in terms of mutation frequency^25^. The glycerol-3-phosphate dehydrogenase (G3PDH) product of *gpsA* plays a role in membrane lipid metabolism and maintenance of intracellular redox potential^26^. We constructed a targeted deletion of *gpsA*, and individually reproduced the identified mutation that generated the G208R variant, in the ancestor D39 strain, using the sweet Janus system^27^.

We assessed the nasopharyngeal colonisation potential of the resulting D39Δ*gpsA* and D39*gpsA*_G208R_ strains relative to a streptomycin resistant D39 control (D39Sm_r_). Both strains showed comparable levels of growth to D39Sm_r_ in nutrient broth (Supplementary Figure 14). However, D39Δ*gpsA* was markedly attenuated in its ability to colonise murine nasopharynx, showing reduced colonisation density relative to D39Sm_r_ from day 1 post infection and evidence of clearance as early as day 3 (Figure 7A). D39*gpsA*_G208R_, by contrast, showed prolonged maintenance of carriage, with 100% of mice still colonised at day 14 post-infection (Figure 7A).

**Figure 7.**
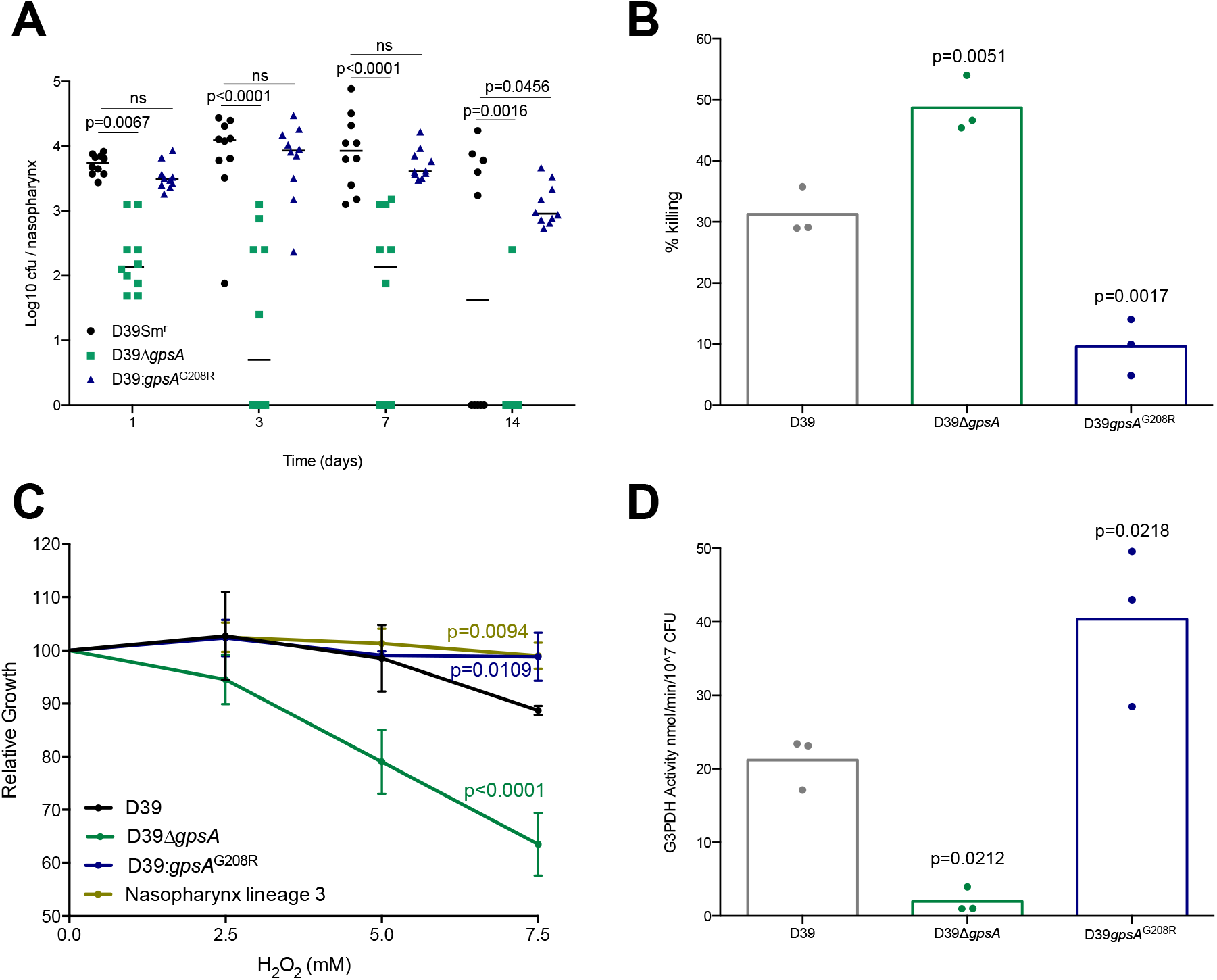
Glycerol-6-phosphate dehydrogenase is required for effective colonisation of nasopharynx. **(A)** Mice were infected with a streptomycin resistant version of the ancestor D39 in which either a clean deletion of *gpsA* (D39Δ*gpsA*) or the SNP causing the G208R substitution (D39:*gpsA*_G208R_) had been generated using the sweet Janus system. Control animals were infected with the streptomycin resistant D39 ancestor expressing wild type *gpsA* (D39Sm_r_ – streptomycin resistant). Infection dose was 1 ×10^5^ colony forming units (cfu) to induce asymptomatic carriage. Bacterial numbers were determined by serial dilution of nasopharynx homogenates onto selective agar at days 1, 3, 7 or 14 post-infection. N = 10 per group. P-values are from two-way ANOVA with Sidak’s multiple comparison test. **(B)** Cfu counts for D39, D39Δ*gpsA* and D39*gpsA*_G208R_following 30 minutes incubation in a 12 mM hydrogen peroxide (H_2_0_2_) solution. Data are presented as percentage killing vs the no H_2_0_2_ control for each strain. P-values are from a one-way ANOVA with Dunnett’s multiple comparison test vs D39. **(C)** Percentage reduction in growth (measured as area under the logistic curve [AUC]) for D39, D39Δ*gpsA*, D39*gpsA*_G208R_ and nasopharynx lineage 3 in the presence of increasing concentrations of H_2_0_2_. P-values are from a two-way ANOVA with Dunnett’s multiple comparison test vs D39. Each point is the mean of three biological replicates, each containing three technical replicates. Error bars are standard deviation. **(D)** Glycerol-3-Phosphate dehydrogenase (G3PDH) enzyme activity measured in the ancestor, D39Δ*gpsA* and D39*gpsA*_G208R_ using a colorimetric assay. Activity was determined by comparison of absorbance readings at 450 nm to a NADH standard curve and are presented as nmol NADH per minute per 10^7^ cfu. P-values are from a one-way ANOVA with Dunnett’s multiple comparison test vs the D39 ancestor. In **(B and D)** each data point represents a single biological replicate, each of which is the mean of three **(B)** or two **(D)** technical replicates.

Because it has been proposed that GpsA is involved in the modulation of intracellular redox potential^26^, we sought to determine whether the G208R substitution in nasopharynx lineages 3 and 4 might impact resistance to reactive oxygen species (ROS). Pneumococci produce ROS as a byproduct of metabolism and are exposed to oxidative stress in the respiratory tract, both from co-colonising microbes and from the actions of neutrophils^28^. A hydrogen peroxide sensitivity assay revealed the significantly elevated susceptibility of D39Δ*gpsA* compared to D39, whereas D39*gpsA*_G208R_ was more resistant to H_2_O_2_ induced killing (Figure 7B). During long-term exposure to H_2_O_2_, both D39*gpsA*_G208R_ and a nasopharynx-passaged lineage, containing the same mutation, showed increased resistance, compared to both D39 and D39Δ*gpsA* (Figure 7C). Analysis of G3PDH activity confirmed a substantial reduction in D39Δ*gpsA* and a significantly elevated level of activity in D39*gpsA*_G208R_, relative to D39 (Figure 7D). These data are consistent with the G208R variant of GpsA increasing the activity of G3PDH, and enhancing the nasopharyngeal colonisation potential of *S. pneumoniae* via increased resistance to oxidative stress.

## Discussion

Experimental evolution is a powerful tool for identification of genetic elements under selection within different environments^4^. Cooper and colleagues recently showed the value of this approach for *Streptococcus pneumoniae* in mouse models, using a nasopharyngeal colonisation model similar to that described here, but using a 19F pneumococcal strain (BHN97x)^29^. There are some interesting commonalities in the findings of the two studies. Firstly, despite the bottlenecks occurring during experimental passage, both studies showed evidence of strong selection operating on pneumococcal populations. Parallelism and convergence were evident in independently evolving lineages within both studies and when comparing lineages derived from one study to those from the other, despite the different strain choice.

Cooper *et al* highlighted the acquisition of frameshift mutations in *mutS* and demonstrated that loss of this gene conferred a hypermutator phenotype. We observed mutations in *mutS, mutS2, mutL* and *mutX*, with those in *mutX* reaching fixation in 2 lineages. Mutations in the *mut* DNA repair genes and in *recN*, which collectively were found in 6 of 20 lineages, might provide an explanation for the increasing numbers of variants we observed per lineage with increasing passage numbers.

Acquisition of mutations in genes encoding components of ABC transport systems was a further feature of both studies. We highlighted the *yknX* mutation in 2 nasopharynx lineages and one lung lineage, but we also observed a mutation in *yknY* that reached fixation in lineage 9. Cooper and colleagues found a mutation in *yknZ* in one nasopharynx-passaged lineage. Collectively, YknXYZ form an ABC transport system involved in resistance to both antibiotics and host-derived antimicrobial peptides^30,31^. The identification of SNPs in the all three genes of this transport system suggests it may play a role in upper-airway colonisation. Further, Cooper *et al* report mutations in *yheS_1, yheS_2* and *yheS_3*, whilst we identified a SNP within *yheH*. These genes also encode ABC transporter components, although the Yhe system is poorly characterised in pneumococcus. Non-synonymous mutations in *yheS_2* were previously identified when comparing paired carriage and disease isolates of serotype 22F^32^, suggesting that this gene may be involved in niche adaptation. Finally, both studies identified mutations in genes with roles in thiamine, copper and pyruvate uptake or metabolism, highlighting the key requirement for trace metal scavenging and regulation of glycolysis during colonisation^33^.

Although experiments were established with D39 stocks that originated from a single colony, we identified some potentially important genetic variation within the population used to found the twenty lineages. Comparison of genetic ancestries of the lineages (Figures 4, 5 and Supplementary Figure 4-13) suggests that the sub-populations of the ancestor bearing the *P0_01974* and *dnaK* mutations were distinct, with each subsequently giving rise to separate genotypes. One of these often drove the other to extinction and, beyond passage 10, most lineages had lost either the *dnaK*-derived genotypes or those arising from the *P0_01974* genotype. In lung lineages, the *dnaK* mutation present in the ancestor became fixed in 8/10 lineages but was lost from the other two, where instead the *P0_01974* mutation became fixed. In nasopharynx lineages, both mutations were eventually lost in 5/10 lineages. Examples of all three outcomes are shown in Supplementary Figure 15. It is possible that, within the nasopharynx, the *dnaK* or *P0_01974* mutations themselves did not significantly affect fitness and that the success of one or the other genotype came down to which was first to acquire a beneficial mutation, or to survive a population bottleneck. Intriguing, however, is the relative success of the *dnaK* mutation in lung lineages, where it became fixed in eight of ten lineages.

Although the lung is not the primary niche of pneumococcus, we opted to experimentally passage D39 through the lung environment, as colonisation of this niche is associated with severe disease in humans. The nasopharynx and lungs differ in several important ways, including temperature, pH, availability of nutrients and degree of immune surveillance^34-36^. Pneumococcal lineages differ in their abilities to cause pneumonia and invasive pneumococcal disease^37^, consistent with some being better adapted to thrive in the lung environment than others. By bypassing the carriage step that would ordinarily precede lung infection we hoped to select for traits specifically advantageous in the lung environment that might, therefore, provide information about how the pneumococcus is able to cause disease. We observed only ten variants that were shared by nasopharynx and lung lineages but were not present in the control passage or ancestor, as compared to 245 variants unique to nasopharynx lineages and 118 unique to lung. Whilst it is likely that only a small fraction of identified variants could be considered adaptive mutations under positive selection, it was notable that the examples of parallelism or convergence we identified tended to be niche-specific. Mutations in *gpsA* and *pvaA* were only observed in nasopharynx lineages, whilst those in *mutX1, lafA* and *sdhB* were unique to lung lineages. The finding that nasopharynx-evolved lineages, but not lung-evolved lineages, showed enhanced carriage potential further suggests that adaptations are at least partially niche-specific. Only *yknX* showed evidence of selection in both niches, with a mutation causing a leucine to valine change at position 333 becoming fixed in one lung lineage, and a SNP causing a serine to asparagine substitution at position 331 becoming fixed in two nasopharynx lineages.

The same non-synonymous mutation in *gpsA* was identified in two nasopharynx lineages and a recent study of pneumococcal carriage in infants in the Gambia also identified a high frequency of mutation in this gene^25^ (Supplementary Table 2). Deletion of *gpsA* in the ancestor strain compromised nasopharyngeal carriage, whilst the identified SNP conferred increased carriage potential and resistance to oxidative stress, when reproduced individually in D39. Glycerol 3-phosphate dehydrogenase reduces NADP^+^ to NADPH during the conversion of sn-glycerol 3-phopshate to glycerine phosphate, thereby helping maintain intracellular redox potential. Increased redox control during carriage may be beneficial in the context of competition with co-colonising microbes or in resistance to neutrophil responses. Mutations conferring such phenotypes may also have consequences for antimicrobial resistance in pneumococcus, as antibiotics including penicillin and ciprofloxacin induce production of ROS^26,38^.

We identified increased adhesion to airway epithelial cells in three of ten nasopharynx-passaged lineages and reduced pneumolysin production in nine of ten lineages. Both these traits could compromise transmission whilst promoting stable carriage. An assessment of transmission potential in niche-adapted lineages will be an important next step for this work. If nasopharynx-passaged lineages have increased potential for non-invasive carriage together with loss of transmission potential, then they may be suitable starting points for the development of live-attenuated pneumococcal vaccines, although requiring further laboratory attenuation to prevent reversion to virulence or acquisition of new traits *in vivo* via recombination.

An open question is the extent to which the genetic changes we observed might be constrained in a natural infection setting. In our model, the process of transmission – associated with tight bottlenecks^16^ – was bypassed by manual transfer of pneumococci between animals. The need to balance attachment and colonisation with shedding and transmission is likely a major determining factor in the trajectory of pneumococcal evolution and the trade-off between transmissibility and within-host fitness has been recently highlighted^39^. The authors outline the role played by competitive exclusion in restricting population diversity, whereby the first variant to colonise prevents later arrivals from doing so. The extent to which this phenomenon might impact evolution in our experimental system, which utilised manual administration of the bacterial population as opposed to natural exposure, is unclear. We did observe loss of some genotypes between passages but also instances where multiple, co-existing genotypes were maintained for several passages.

One important caveat to the findings presented relates to the anatomical, biochemical and immunological differences between the mouse and human respiratory tracts. In addition to adapting pneumococci to the nasopharynx or to the lungs, the passage process we describe might select for variants that give an advantage specific to colonisation of murine respiratory tissue. We are, however, encouraged by a recent study in humans, analysing microevolution of pneumococci during natural colonisation of infants^25^. Twenty (43%) of the genes in which we observed fixed non-synonymous SNPs in nasopharynx-passaged lineages were also targets of mutation during prolonged colonisation in infants (Supplementary Table 1). These included *gpsA* and *yknX*, in which we observed parallel evolution. Considering all the genes in which we identified gene variants (synonymous or non-synonymous SNPs or indels) in nasopharynx passaged lineages, 39% (68/174) were also identified in the dataset of Chaguza and colleagues (Supplementary Dataset 2).

Collectively, our findings suggest that *in vivo* experimental evolution can be a valuable tool for the study of host-pathogen interactions and disease mechanisms. The local environment shapes pneumococcal evolution, and differences between the niches of the upper and lower airway are sufficient to alter the trajectory of evolution in populations of bacteria exposed to one or the other. We identified several examples of parallel evolution in independently passaged populations and the genes under selection highlight the importance of cell surface modification and metabolic versatility in pneumococcal colonisation and disease processes. *In vivo* experimental evolution approaches can be used to identify pathogen genes with roles in within-host processes, as exemplified by *gpsA*, that appeared to be under strong selection in nasopharynx and in which we identified a single nucleotide change conferring phenotypic change and increased within-host fitness.

## Methods

### Media and growth conditions

*S. pneumoniae* D39 (serotype 2, NCTC 7466) isolates were grown at 37°C in 5% CO_2_ on blood agar base (BAB) (Oxoid) supplemented with 5% (vol/vol) defibrinated horse blood (Oxoid) and 1 µg/ml of gentamycin and overnight cultures were prepared in 10 ml of brain heart infusion (BHI) broth (Sigma). Bacteria were identified as pneumococci by α-hemolysis on blood agar and by optochin sensitivity. The D39 used in this study was generously provided by Prof Tim Mitchell (University of Birmingham, UK).

### Ethics statement

All mouse infection work was performed at the University of Liverpool with prior approval by the UK Home Office (project licence PB6DE83DA) and the University of Liverpool Ethics Committee. For all experiments, 6 to 8 week old female CD1 mice were purchased from Charles River (Margate, UK). Mice were randomly assigned to a cage of 1-5 mice, depending on the experiment, on arrival at the unit, by staff with no role in study design. Mice were housed in individually ventilated cages and allowed to acclimatise for 7 days prior to infection.

### Experimental evolution in mouse infection models

A standardised suspension of the D39 ancestor strain was prepared for inoculation by overnight growth in BHI from a single colony. The next day, sub-culture was performed and the bacteria allowed to grow to mid-exponential phase (∼6 hours) before storage at −80°C. For the initial infection, this suspension was thawed at room temperature and bacteria were harvested by centrifugation and suspended in PBS. For the carriage model, 10 mice were intranasally infected under light anaesthesia, using a mix of oxygen and isoflurane, with 1 × 10^5^ colony forming units (cfu) in 10 µl saline. To induce pneumonia, 10 mice were infected intranasally with 1.5 × 10^6^ cfu, in 50 µl saline, to ensure pneumococci reached and colonised the lung. The 20 infected mice were treated as founders of 20 independent pneumococcal lineages, 10 of which were passaged through nasopharynx and 10 which were passaged through lung.

In the carriage model, the bacteria remained in the nasopharynx for 7 days for each passage and, for the pneumonia model, bacteria were in the lung environment for 48hrs, or less if the mice reached the severity endpoint before this time. The time the lineages spent in the lungs ranged from 22-48 hours per passage, with an average time of 31.91 hours. After each passage, pneumococci were recovered from the nasopharynx, in the carriage model, or lungs, in the pneumonia model. After minimal passage on gentamicin BAB agar with an optochin disk to confirm identity as *S. pneumoniae*, the entire population of recovered pneumococci for each lineage was resuspended into 1 ml of BHI/10% glycerol, which was split 500:250:250 between three tubes and stored at −80°C for future use in infection experiments, phenotypic analysis or genome sequencing. One of the 250 µl frozen aliquot tubes was used to determine the cfu by Miles and Misra and to re-infect a new mouse for the next round of passage. This passage process was repeated a total of 20 times for each of the separately evolving lineages in the carriage and pneumonia models of infection. Figure 1 gives an overview of the *in vivo* experimental evolution process undertaken during this study.

### Control laboratory passage on BAB agar

The D39 ancestor freezer stock used for the initial infection in the *in vivo* evolution experiments was thawed and used to streak 10 gentamicin BAB plates with an optochin disc and incubated for 20-24 hours at 37°C in 5%. For the next passage for each lineage, the population was re-streaked onto a fresh gentamicin BAB plate and this process was repeated 20 times for the 10 separately evolving lineages. Freezer stocks were made from passages 1, 5, 10, 15 and 20 and stored for future analysis. In the final analysis, only 8 of the 10 lineages were included due to contamination of the DNA used for sequencing.

### Illumina sequencing and read processing

DNA extraction protocols are detailed in supplementary methods. Short read sequencing was performed to produce 151-bp, paired-end reads on an Illumina HiSeq2500 at the Wellcome Trust Sanger Institute, UK. This was carried out for the ancestor strain and for passages 1, 5, 10, 15 and 20, for both the nasopharynx and lung evolved lineages. Short read sequencing was also carried out for the passage 20 control lineages. To obtain a deep sequence coverage of the ancestor D39 strain, DNA was paired-end sequenced on an Illumina NextSeq 500 system using 150-bp read lengths, by Vertis Biotechnologie AG, Germany. All bioinformatic analysis of sequencing data was carried out using a virtual machine hosted by the Cloud Infrastructure for Microbial Bioinformatics (CLIMB) consortium^40^. Raw Illumina reads were processed using Trim Galore (v0.4.4) with Cutadapt (v1.9.1) for paired-end reads, using default settings, to remove short low quality reads (< 20 bp), trim Illumina adaptor sequences, and eliminate poor-quality bases from the sequences (< Q20). FastQC (v0.11.5) confirmed that the resulting trimmed reads were of sufficient quality for subsequent analysis.

### PacBio long read sequencing and genome assembly

For the ancestor strain, single molecule real-time (SMRT) sequencing (Pacific Biosciences, PacBio, CA, USA) of the ancestor D39 isolate was performed at the Wellcome Trust Sanger Institute (Cambridge, UK). The resulting BAM file was converted to FASTQ with bamToFastq (v2.28.0). PacBio long reads were filtered by quality using Filtlong (v0.2.0-https://github.com/rrwick/Filtlong) and assembled with the Flye Assembler (v2.4.2)^41^; default settings were applied for both programmes. The resulting contig, with 469x coverage depth, was polished using the PacBio SMRT minimap2 tool (v2.15-r905) and Racon Polisher (v1.3.3-https://github.com/isovic/racon), with default settings applied. A consensus sequence of the ancestor was generated using PacBio SMRT tools Blasr alignment (v5.1) and Arrow (v2.3.3). This was then corrected by aligning the sequence with the ancestor Illumina reads using bwa mem (v0.7.17-r1188) and re-assembling with Pilon (v1.22)^42^. The resulting corrected consensus sequence was annotated using Prokka (v1.13)^19^ and run through the Quality Assessment Tool for Genome Assemblies (QUAST, v4.6.3) to obtain the assembly statistics. Comparison of Prokka annotation with the recent D39 genome assembly of Slager *et al*.^18^ is detailed in supplementary methods.

### Variant calling

Variants were detected from the processed Illumina reads for the nasopharynx and lung evolved lineages at passages 1, 5, 10, 15 and 20, as well as the control lineages at passage 20. The short sequence reads were mapped to the D39 ancestor corrected consensus sequence to identify mutations, using the Breseq programme (v0.31.1)^43^. Default settings were applied with the predict-polymorphisms function. This process identified the percentage frequency of variants within a lineage population at each passage. Synonymous and non-synonymous single nucleotide polymorphisms (SNPs) as well as insertions/deletions and intergenic mutations were detected. The deeper sequenced lllumina reads for the ancestor strain (x300 coverage) were also used to detect variants within the original D39 ancestor inoculum, with the polymorphism-frequency-cut off parameter set to >2%.

### Comparison with clinical isolate data

We analysed pneumococcal genomes from a recently published dataset of isolates sampled longitudinally in newborn infants, to assess whether the mutations identified during experimental evolution occur frequently within hosts during natural carriage episodes^25^. We mapped the reads of each isolate against the ancestral D39 reference genome and generated a consensus alignment for each episode using Snippy (v4.3.6) (https://github.com/tseemann/snippy). The genomic positions with SNPs during each carriage episode were identified, annotated, and compared between episodes to identify episodes with changes occurring in parallel at the same location using BioPython^44^.

### Survival and time point pneumonia experiments

Mice were infected with standardised pneumococcal isolate stocks at a dose of 1 × 10^6^ cfu in 50 µl saline via intranasal administration under light anaesthesia, to induce acute pneumococcal disease. Mice were monitored for signs of disease and pain score was determined using the scheme of Morton^45^. For survival experiments, mice were monitored for 7 days and culled once lethargic. For time point experiments, mice were culled at 0, 12, 24 and 36 hours post-infection or if they reached the experimental endpoint (lethargy). Lungs and blood samples were taken and pneumococcal cfu determined via Miles and Misra dilution onto gentamycin BAB plates. Construction of SNP and deletion mutants for use in these experiments is described in Supplementary Methods.

### Carriage time point experiments

Mice were infected as described above but with a dose of 1 × 10^5^ cfu in 10 µl saline to induce asymptomatic nasopharyngeal colonisation. At 1, 3, 7, 14 and 21 days post infection, mice were culled, nasopharynx samples taken and pneumococcal cfu determined.

### Analysing pneumococcal growth dynamics in the presence of hydrogen peroxide

Growth assessment was performed as detailed in Supplementary Methods, with the following modifications. Pneumococcal isolates were grown in triplicate in BHI in 96-well plates in the presence of hydrogen peroxide (H_2_0_2_, Sigma) at concentrations 0, 2.5, 5.0 and 7.5 mM. Growth dynamics were analysed with the R package GrowthCurver and the percentage reduction in the AUC values for each isolate in the presence of H_2_0_2_ was calculated in comparison to 0 mM.

### Hydrogen peroxide sensitivity assays

Mid-log phase pneumococci were added in duplicate to microtiter plate wells and incubated at 37°C for 30 minutes in BHI with 12 mM of H_2_0_2_. The CFU/mL of each isolate following the incubation was determined via Miles and Misra dilution onto gentamycin BAB plates and the percentage killing was calculated for each test isolate in comparison to the untreated (BHI only) control.

### Glycerol-3-phosphate dehydrogenase assay

G3PDH activity was measured using a colorimetric assay kit (Abcam). Overnight cultures of isolates were centrifuged for 15 mins at 3000 rpm and the pellet resuspended in 200 ul of G3PDH assay buffer and lysed with the addition of sodium deoxycholate at 10%. This was incubated at room temperature for 10 minutes, centrifuged for 15 minutes at 3000 rpm and the supernatant used for the assay following the manufacturer’s instructions. Absorbance readings at 450 nm were taken at time zero and following a 60 minute incubation at 37°C and the activity of G3PDH for each isolate was calculated by comparison to a NADH standard curve.

### Statistical analysis

GraphPad Prism version 8.2.1 was used for statistical analysis. Statistical tests undertaken for individual experiments are detailed in the respective figure legends. p<0.05 was considered to be statistically significant. Data were tested for normality and to define the variance of each group tested. All multi-parameter analyses included corrections for multiple comparisons and data are presented as mean±standard deviation (SD) unless otherwise stated.

## Author contributions

DN conceived the study. AG, JH, SB, JR and DN designed experiments. AG, FL, DH, CM, HE and DN performed experiments. AG, CC, FL, TB, JH, JR and DN analysed data. SB, JR and DN provided resources and reagents and supervised staff. AG and DN wrote the manuscript, with input from all authors.

## Acknowledgments

DN and AG are supported by a Sir Henry Dale Fellowship, awarded to DN and funded by the Wellcome Trust and the Royal Society (Grant number 204457/Z/16/Z). FL is supported by a PhD studentship from the Rosetrees Trust. TB was supported by a Harry Smith Vacation Studentship awarded by the Microbiology Society (grant number GA002322). JCDH is funded be a Wellcome Trust Senior Investigator Award (106914/Z/15/Z). JR is supported by 1U01AI124302 and 1RO1AI110618, and by ALSAC. We would like to thank Siobhan Austin-Guest and Katherine Auger at the Wellcome Sanger institute for facilitating sequencing, and the Veening lab, Université de Lausanne, for creation and maintenance of Pneumobrowse, which has been an invaluable resource throughout this study.

## Notes

### Competing Interest Statement

The authors have declared no competing interest.

