## Supplementary Figures and Methods for "Identification of colonisation and virulence determinants of *Streptococcus pneumoniae* via experimental evolution in mouse infection models"

Angharad E Green et al.

### Supplementary Methods

#### DNA extractions

Genomic DNA was extracted from fresh overnight cultures of pneumococcal isolates. A pre-lysis step was included where a pellet from 2 ml of overnight culture was incubated for 1 hour at 37$_{o}$C with lysozyme (Sigma) and mutanolysin (Sigma) in TE buffer. Proteinase K (Qiagen) and Buffer AL (Qiagen) were then added and samples incubated at 56$_{o}$C for 2 hours. The DNeasy Blood & Tissue Kit (Qiagen) was then used, following the manufacturer's instructions for purification of DNA from Gram-positive bacteria. Quality assessment of DNA was carried out by measuring the absorbance ratios 260/280 nm and 260/230 nm using a NanoDrop Spectrophotometer (Thermo Fisher Scientific); ratios of ≥1.8 were used as an indication of good quality DNA. A Qubit™ fluorometer system with the broad range DNA kit (Invitrogen) was used to quantify DNA, according to manufacturer's instructions. We required 10 µg of DNA at 200 ng/µl in a final volume of 50 µl for PacBio sequencing and for Illumina sequencing >1 µg of DNA between 20-150 ng/µl in a final volume of 25 µl was needed.

#### BLAST comparison of the D39 ancestor strain sequence annotation

In order to check for annotations of those genes labelled as hypothetical proteins in the Prokka (v1.13) annotated D39 ancestor genome, we compared all genes to those in the recently published assembly of the D39 strain[1], using the command line standalone NCBI Nucleotide-Nucelotide BLAST programme (BLASTp, v2.9.0+)[2]. Matches between genes in the two strains were determined by applying an E-value cut-off of 1E-40, and setting the best-hit filtering algorithm overhang parameter to 0.25. The results can be found in Supplementary Dataset 1.

**Confirmation of SNPs in niche evolved lineages**

DNA was extracted from the ancestor D39 strain and the lineages of interest as previously described. To confirm SNPs, the target gene was amplified via PCR, in a Veriti 96-well thermal cycler (Applied Biosystems), using the TaKaRa Ex Taq® DNA Polymerase kit (TakaraBio, Germany) with the PCR primers described in Supplementary Dataset 3 (primers were purchased from Eurofins Genomics). To confirm amplification of the relevant gene, PCR products were run using a 1% agarose gel in TAE buffer (Invitrogen). To extract and purify the PCR products from the gel, the MinElute Gel Extraction Kit (Qiagen) was used following manufacturer's instructions. The purified PCR products were quantified using the Nanodrop and sent for sequencing at Eurofins Genomics via the TubeSeq service. The resulting forward and reverse sequences of were aligned using the MEGA alignment software (version 10.1.5) against the relevant gene sequences to confirm the presence of the SNPs in the niche evolved lineages and the SNP absence in the D39 ancestor strain. All primer sequences can be found in Supplementary Dataset 3.

**Construction of Muller plots, genotype frequency plots and genotype ancestry trees**

The command line programme muller (v0.6.0 - https://pypi.org/project/muller/) with default parameters applied, was used to produce the genotypes and trajectories tables for each of the 20 nasopharynx and lung evolved lineages. These tables were then were used as inputs for ggplot2 (v3.2.0) and ggmuller (v0.5.3) in R-Studio (v3.6.1), to produce Muller plots, genotype frequency plots and genotype ancestry trees for each lineage. Genes which had variants in two or more lineages were manually coloured the same across the different plots. The genotypes found in the ancestral strain were coloured as white in each lineage Muller plot, except where Breseq assigned the *dnaK* mutation or *P0_01974* ancestral SNPs into other genotypes.

**Cluster of Orthologous Group (COG) functional annotation**

The COG functional annotation[3] of the D39 Prokka predicted CDS was carried out using the online eggNOG-mapper (v2) database using default settings[4,5]. The assigned COG categories of genes with all types of mutations found in the lung and nasopharynx passage 20 isolates were analysed to determine which COG functional groups were represented amongst the variant genes. The COG categories for those genes with mutations which appear in more than one lineage, are represented only once and gene variants also found in the control passage were removed from this analysis. The percentage of COG functional groups were determined and the analysis was represented as pie charts. The percentages are based on the total number of variant genes in the lung and nasopharynx passage 20 lineages.

**Visualising the non-synonymous mutations on the D39 ancestor genome**

The Circa software (OMGenomics, http://omgenomics.com/circa/) was used to construct Circos plots for visualisation of lineage SNPs on the D39 ancestor genome. The Prokka generated GFF file was converted to a tab-separated format that indicated the start and end positions and names of the 2069 D39 ancestor genes, these are displayed as grey lines on the 1st track of the genome maps. The positions of the genes containing non-synonymous SNPs, identified in twenty-times passaged pneumococci from the lung and nasopharynx, were drawn on the maps with circles.

**Mutant construction**

The D39 ancestor strain was first made resistant to streptomycin by introducing a point mutation in the *rpsL* gene to produce a K56T variant. A SOEing PCR using the TaKaRa Ex Taq® DNA Polymerase kit (TakaraBio, Germany) was performed with the streptomycin resistant up_fragment and down_fragment primers (forward and reverse) (all primer sequences can be found in Supplementary Dataset 3). The D39 ancestor strain was then transformed using the resulting SOEing PCR product by growing the strain at 37°C with 5% $CO_2$ in C+Y media. Once an OD of 0.07-0.1 (500 nm) was reached, 1 ml was transferred to a 1.5 ml tube and 3 µl competence stimulating peptide (CSP)1 was added along with the PCR product. This was incubated for 3 hours at 37°C with 5% $CO_2$. Mutants were selected on TSA + 3% blood + 20 ug/mL neomycin + 800 ug/mL streptomycin. The resulting streptomycin resistant strain was then used for subsequent mutant construction.

Targeted deletion of the *gpsA* (D39Δ*gpsA*) was carried out using the sweet Janus system[6]. The sweet Janus cassette was amplified using the SweetJanus forward and reverse primers. The flanking regions upstream and downstream of the *gpsA* locus (~1 kb) were amplified to produce the PCR products GpsA_Up_SJ and GpsA_Down_SJ. A SOEing PCR product was produced by combining both the flanking region and the Sweet Janus PCR products. To delete *gpsA*, the resulting SOEing PCR product was used to transform the D39 streptomycin resistant ancestor strain described above. The mutants were selected by plating 200 µl of transformation product on TSA + 3% blood + 20 µg/mL neomycin + 400 µg/mL kanamycin. Selected mutants were also plated onto TSA + 3% blood + 20 µg/mL neomycin + 800 µg/mL streptomycin + 10% sucrose to confirm loss of streptomycin resistance.

To create the G208R point mutation in the *gpsA* gene, the desired gene region was amplified to swap into the locus site previously deleted in D39Δ*gpsA*. The PCR parts

GpsA_G208R_Up and GpsA_G208R_Down were produced using combinations of the R primer (for upstream) and F primer (for downstream) with overlapping sequences of the *gpsA* point mutation (Supplementary Dataset 3). A SOEing PCR product was produced by combining these PCR parts and the resulting product was used to transform D39Δ*gpsA* as previously described. The *gpsA* point mutation mutants were selected on TSA + 3% blood + 20 µg/mL neomycin + 800 µg/mL streptomycin + 10% sucrose, no growth on kanamycin was also verified. To confirm successful transformation of the knockout and the point mutation mutants, primers outside of the transformed regions were used for PCR and the region was subsequently sequenced.

**Analysing growth dynamics of pneumococcal isolates**

Pneumococcal isolates were streaked onto gentamycin BAB plates and incubated overnight at 37°C in 5% $CO_2$. A loopful of colonies were resuspended into BHI and 200 µl of culture with a starting OD of approximately 0.002 at 500 nm, was added to the relevant wells of a 96 well plate (Corning Laboratories, Corning, NY). BHI broth was used as a negative control to make sure there was no contamination. The plate was incubated in a Varioskan multimode microplate reader (TheremoFisher Scientific) at 37 ˚C with 5% $CO_2$ for 24 hours, taking reads at 500 nm every 15 mins with double orbital shaking for 10 seconds before the reading. The mean OD readings from the negative control wells were subtracted from the test wells and the resulting OD readings were analysed using the R package GrowthCurver[7] (version 0.3.0) in R-Studio (v3.6.1) and isolate growth dynamics were determined and represented as the area under the logistic curve (AUC). This was protocol was repeated with freshly streaked plates to obtain biological replicates of the data.

**Cell lines**

Human lung adenocarcinoma epithelial cell line A549 (ATCC CCL-185) was used to assess adhesion and invasion potential of pneumococci. All cells were tested for mycoplasma contamination prior to starting experiments and confirmed as negative. A549 cells were maintained in Dulbecco's Modified Eagle Medium (DMEM) supplemented with 10% Foetal Bovine Serum (FBS).

**Adhesion/Invasion assays**

Overnight cultures of pneumococci were diluted to an OD of 0.4 at 500 nm and centrifuged at 4000 rpm for 10 min. The pellet was resuspended in DPBS (pH 7.4) and diluted in DMEM supplemented with 5% FBS, for infection of A549 airway epithelial cell monolayers at a multiplicity of infection (MOI) of 10. The plates were centrifuged at 800 rpm for 5 min to synchronise infection and incubated at 37ºC for 1 hour. Following the 1 hour infection, the media was removed from the wells which were then washed 4X with DPBS. After washing, in the wells of the plate designated for assessment of adhesion, 100 µl/well of Trypsin + 0.025% EDTA was added and incubated for 5-10 minutes at 37°C to detach the cells. The cells were then lysed by adding 1 ml/well of 0.025% Triton X-100 diluted in DMEM+FCS and incubated for 10 minutes at 37°C. To determine the cfu/ml of bacteria which had adhered to the A549 cells, samples were serially diluted 1:10 in PBS and 60 µL was plated on gentamycin BAB plates and incubated 37°C in 5% $CO_2$.

For the plates designated invasion, after washing, the monolayers were incubated with DMEM+FBS containing penicillin (10 µg/ml) and gentamicin (400 µg/ml) for 2 h to kill extracellular pneumococci. Following this incubation, cells were lysed and bacteria enumerated as described above. The percentage of pneumococci that adhered/invaded was calculated as (CFU in the lysate / CFU used for infection)×100.

**Pneumolysin Detection ELISA**

Ninety-six-well ELISA microplates (Corning Laboratories, Corning, NY) were coated overnight at 4°C with 1 μg in 100 μl per well mouse anti-Ply (PLY-4) antibody (Abcam, ab71810). After washing 5 times with 300 μl/well of wash buffer (PBS + 0.05% tween), plates were blocked for 3 hours with 200 μl/well PBS + 20% FBS. Pneumococci were prepared from overnight cultures that were centrifuged for 15 mins at 3000 rpm and the pellet resuspended in 10 ml PBS and lysed with the addition of 100 μl of sodium deoxycholate at 10%. This was incubated at room temperature for 10 minutes, centrifuged for 15 minutes at 3000 rpm and the supernatant used for the assay. For the standards, a ten 2-fold pneumolysin toxin dilution series was prepared. After washing the plate, the bacterial lysates and standards at 100 μl/well were added and incubated at room temperature for 2 hours. After another wash, 1 μg/well rabbit anti-Ply antibody (Abcam, ab71811) in 100 μl of diluent was added and incubated for 2 hours. Plates were washed, and goat anti-rabbit–alkaline phosphatase antibody (Abcam, ab6722) was added for 30 minutes at room temperature. After washing, 100 μl/well pNPP colour reagent (Sigma) was added for 30 minutes and incubated in the dark before the reaction was stopped with 100 μl of 1m NaOH. The absorbance was measured at 405 nm in a Varioskan multimode microplate reader (TheremoFisher Scientific).

1 Slager, J., Aprianto, R. & Veening, J. W. Deep genome annotation of the opportunistic human pathogen Streptococcus pneumoniae D39. *Nucleic Acids Res* **46**, 9971-9989, doi:10.1093/nar/gky725 (2018).

2 Camacho, C. *et al.* BLAST+: architecture and applications. *BMC Bioinformatics* **10**, 421, doi:10.1186/1471-2105-10-421 (2009).

3 Tatusov, R. L., Galperin, M. Y., Natale, D. A. & Koonin, E. V. The COG database: a tool for genome-scale analysis of protein functions and evolution. *Nucleic Acids Res* **28**, 33-36, doi:10.1093/nar/28.1.33 (2000).

4 Huerta-Cepas, J. *et al.* Fast Genome-Wide Functional Annotation through Orthology Assignment by eggNOG-Mapper. *Mol Biol Evol* **34**, 2115-2122, doi:10.1093/molbev/msx148 (2017).

5 Huerta-Cepas, J. *et al.* eggNOG 5.0: a hierarchical, functionally and phylogenetically annotated orthology resource based on 5090 organisms and 2502 viruses. *Nucleic Acids Res* **47**, D309-D314, doi:10.1093/nar/gky1085 (2019).

6 Li, Y., Thompson, C. M. & Lipsitch, M. A modified Janus cassette (Sweet Janus) to improve allelic replacement efficiency by high-stringency negative selection in Streptococcus pneumoniae. *PLoS One* **9**, e100510, doi:10.1371/journal.pone.0100510 (2014).

7 Sprouffske, K. & Wagner, A. Growthcurver: an R package for obtaining interpretable metrics from microbial growth curves. *BMC Bioinformatics* **17**, 172, doi:10.1186/s12859-016-1016-7 (2016).

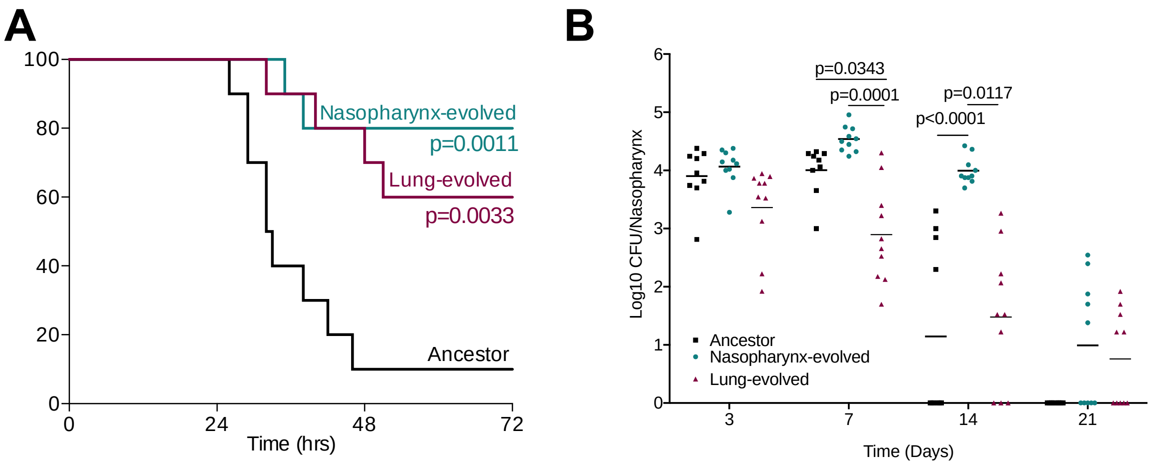


**Supplementary Figure 1. Niche-passaged lineages in pneumonia and nasopharyngeal carriage models. (A)** Survival analysis of mice infection with a pneumonia-inducing dose of ancestor (D39) or twenty times passaged populations from nasopharynx or lungs (n=10 per group). P values are from log-rank (Mantel-Cox) test vs the ancestor. **(B)** Twenty-one day nasopharyngeal carriage of ancestor, or twenty times passaged populations from nasopharynx or lungs (n=10 per group per timepoint). P values are from two-way ANOVA with Sidak's multiple comparisons test.

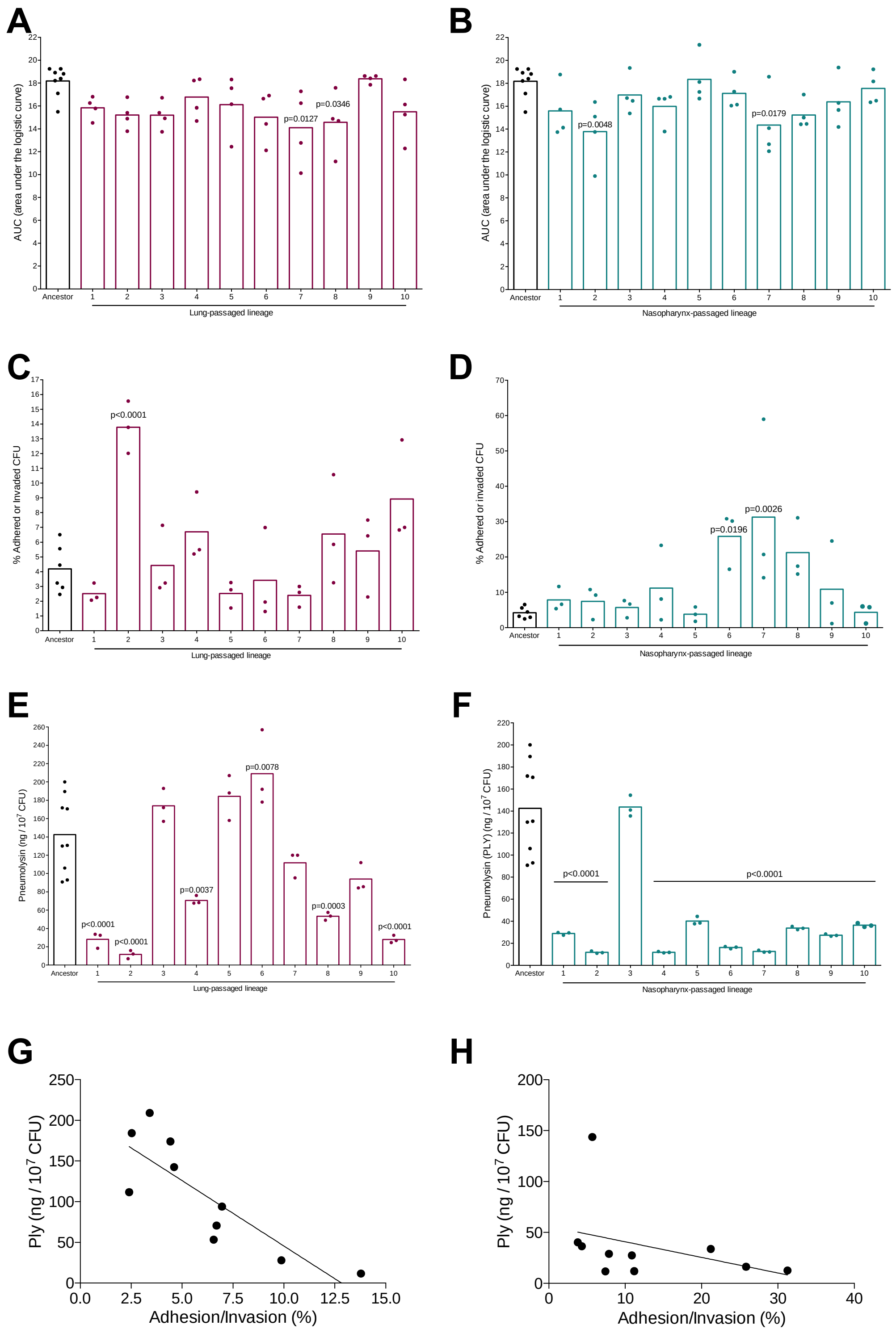


**Supplementary Figure 2. Phenotypes of niche-passaged pneumococci.** Twenty times passaged pneumococcal lineages from lung or nasopharynx were compared to the ancestral D39 population. Growth over 24 hours in nutrient broth is presented as area under the curve (AUC) values for **(A)** lung-passaged and **(B)** nasopharynx-passaged lineages. **(C, D)** Adhesion efficiency to A549 airway epithelial cells was determined following a 1 hour co-incubation. **(E, F)** Pneumolysin concentrations in lysates prepared from 1x10<sub>7</sub> mid-log phase pneumococci were determined by ELISA and comparison to a recombinant pneumolysin standard curve. In **(A-F)** p-values were determined by one-way ANOVA with Dunnett's multiple comparison test vs the ancestor. Data points show individual biological replicates, each of which is the mean of three technical replicates. **(G, H)** For each lineage, mean adhesion percentage was plotted against mean pneumolysin concentration and a line of best fit determined by linear regression.

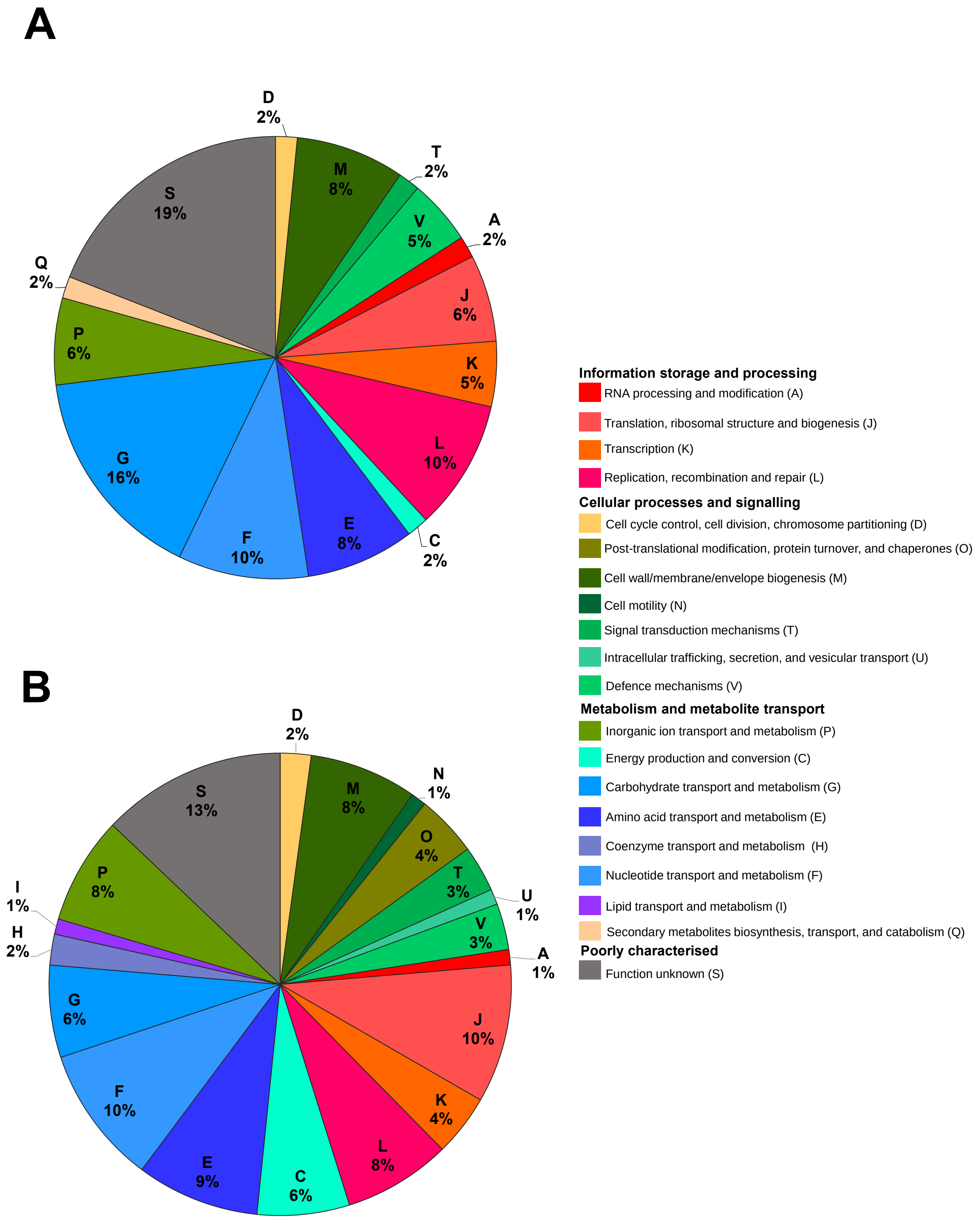


**Supplementary Figure 3. COG analysis of passage 20 variants.** Analysis of variants in (A) lung-passaged, and (B) nasopharynx-passaged lineages. COG categories were assigned to genes for all mutations found in the lung and nasopharynx passage 20 isolates. The percentage of gene variants mapping to the various COG functional groups is represented as pie charts. The COG categories for those genes with mutations appearing in more than one lineage are represented only once and gene variants also found in the control passage were removed from this analysis. The segments of the pie charts are coloured according to the COG-assigned function, as depicted in the key.

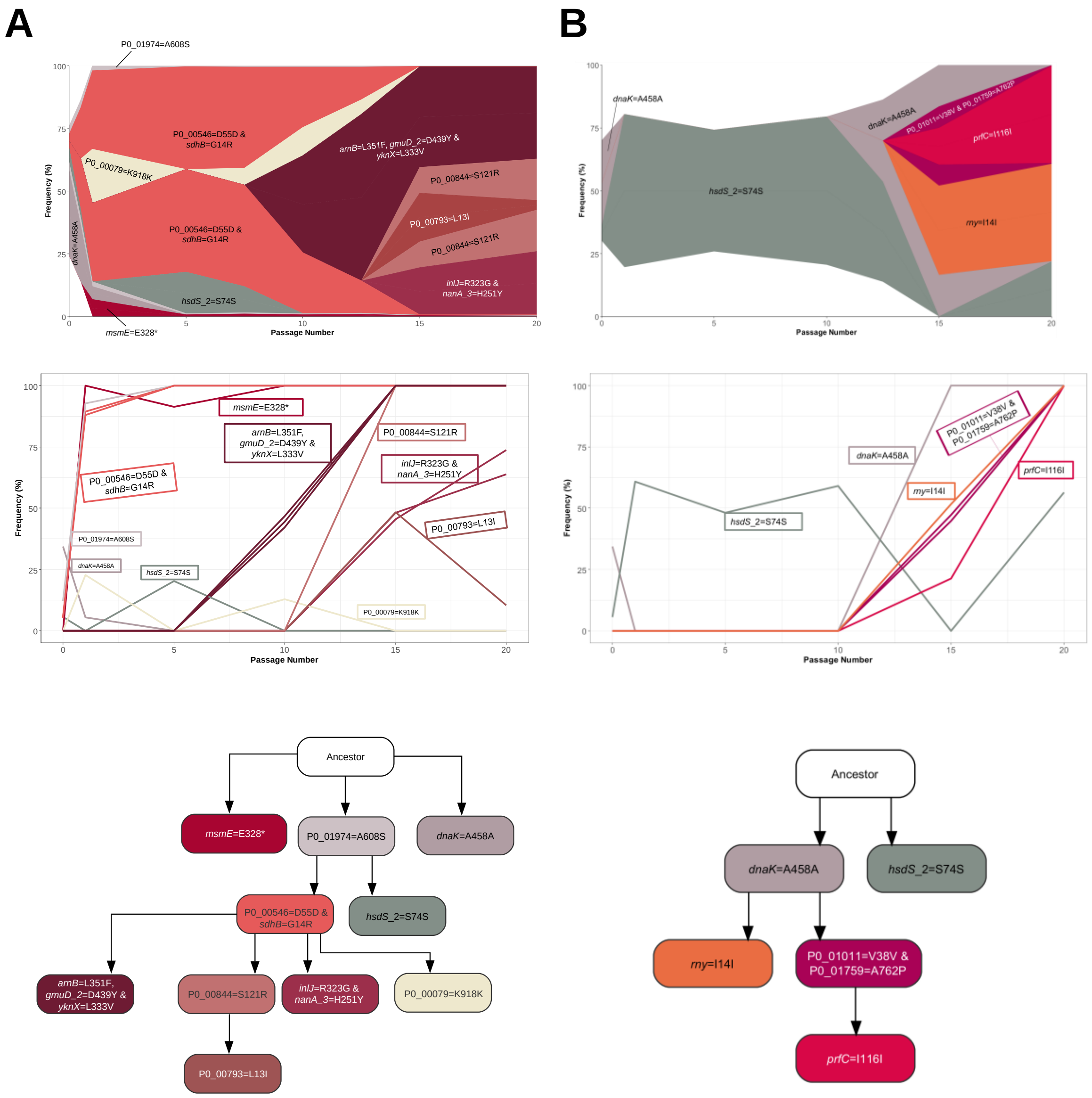


**Supplementary Figure 4. Genotype emergence and dynamics in lung passaged pneumococcal populations. (A)** Lung-passaged lineage 1. **(B)** Lung-passaged lineage 2. Panels, from top-to-bottom, are Muller plots, genotype frequency plots and genotype ancestry trees.

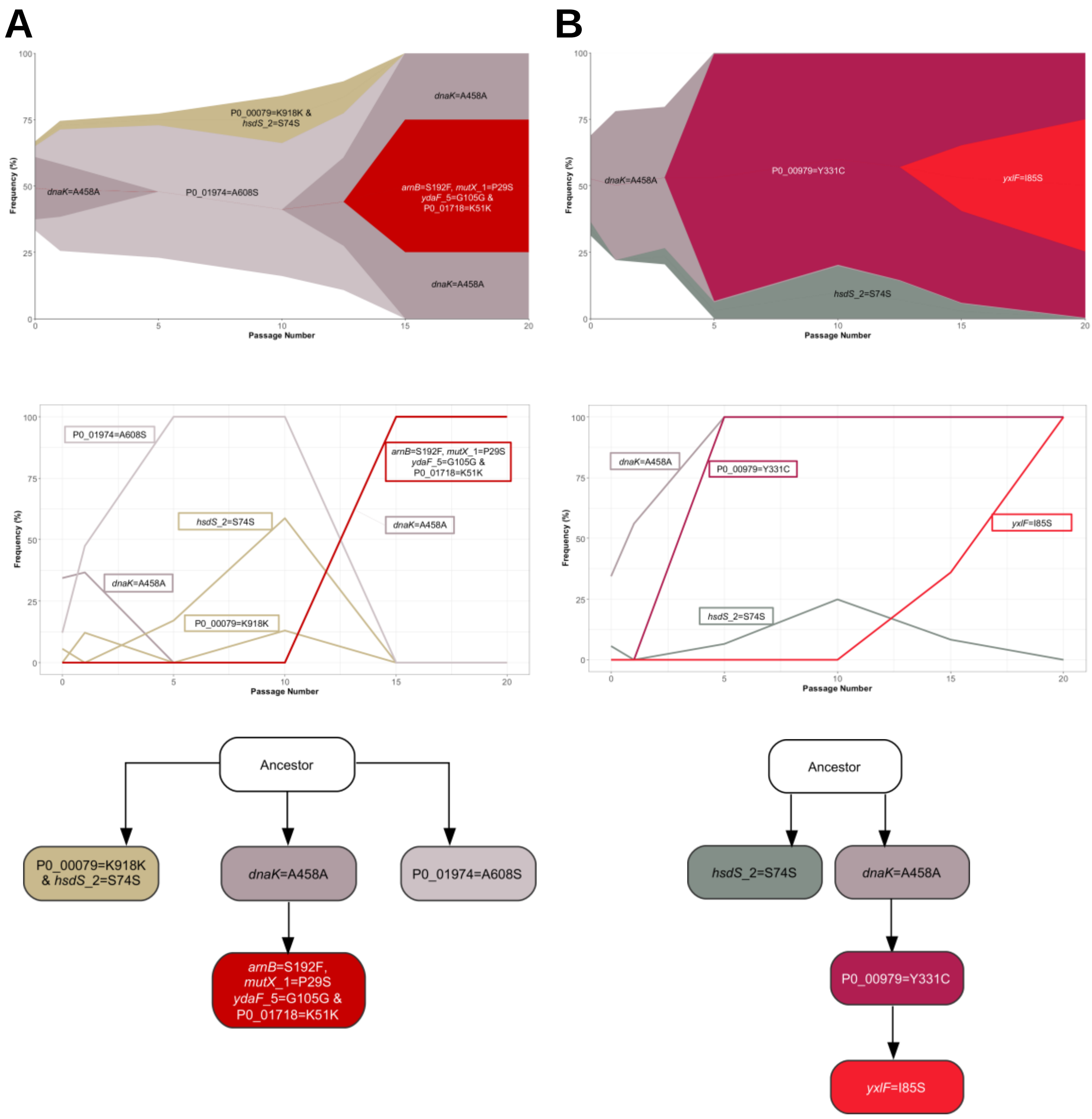


**Supplementary Figure 5. Genotype emergence and dynamics in lung passaged pneumococcal populations. (A)** Lung-passaged lineage 3. **(B)** Lung-passaged lineage 4. Panels, from top-to-bottom, are Muller plots, genotype frequency plots and genotype ancestry trees.

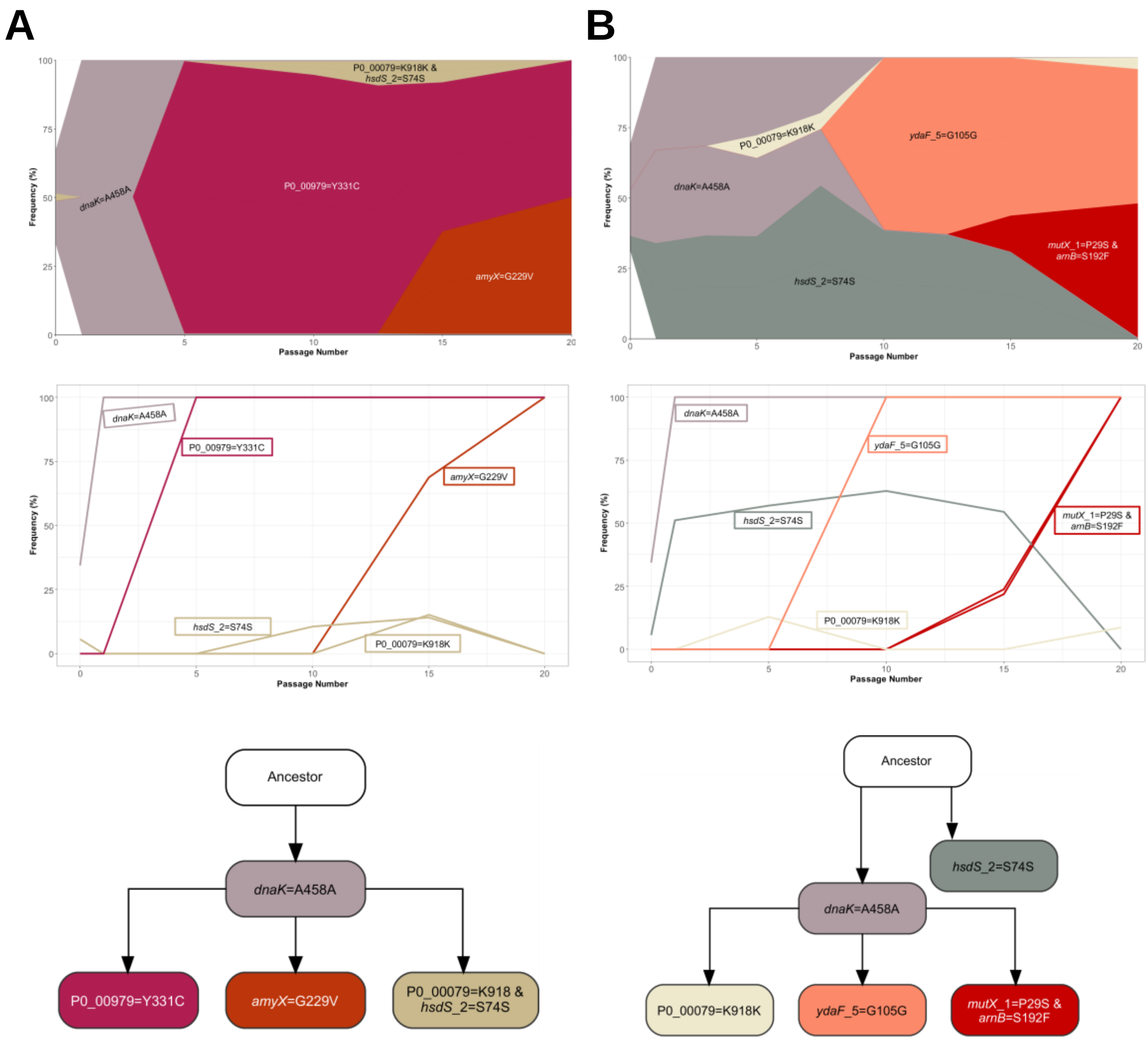


**Supplementary Figure 6. Genotype emergence and dynamics in lung passaged pneumococcal populations. (A)** Lung-passaged lineage 5. **(B)** Lung-passaged lineage 6. Panels, from top-to-bottom, are Muller plots, genotype frequency plots and genotype ancestry trees.

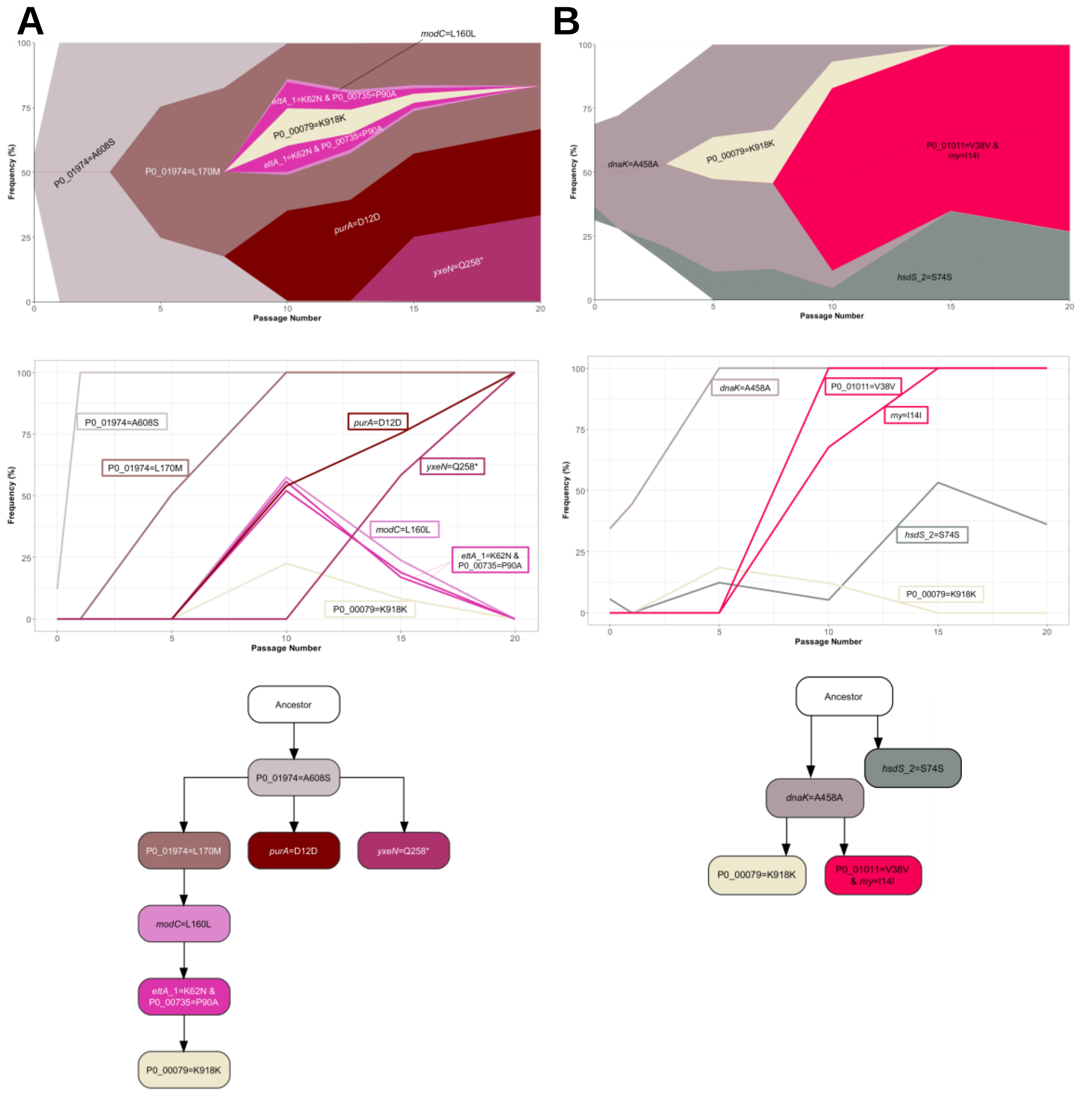


**Supplementary Figure 7. Genotype emergence and dynamics in lung passaged pneumococcal populations. (A)** Lung-passaged lineage 7. **(B)** Lung-passaged lineage 8. Panels, from top-to-bottom, are Muller plots, genotype frequency plots and genotype ancestry trees.

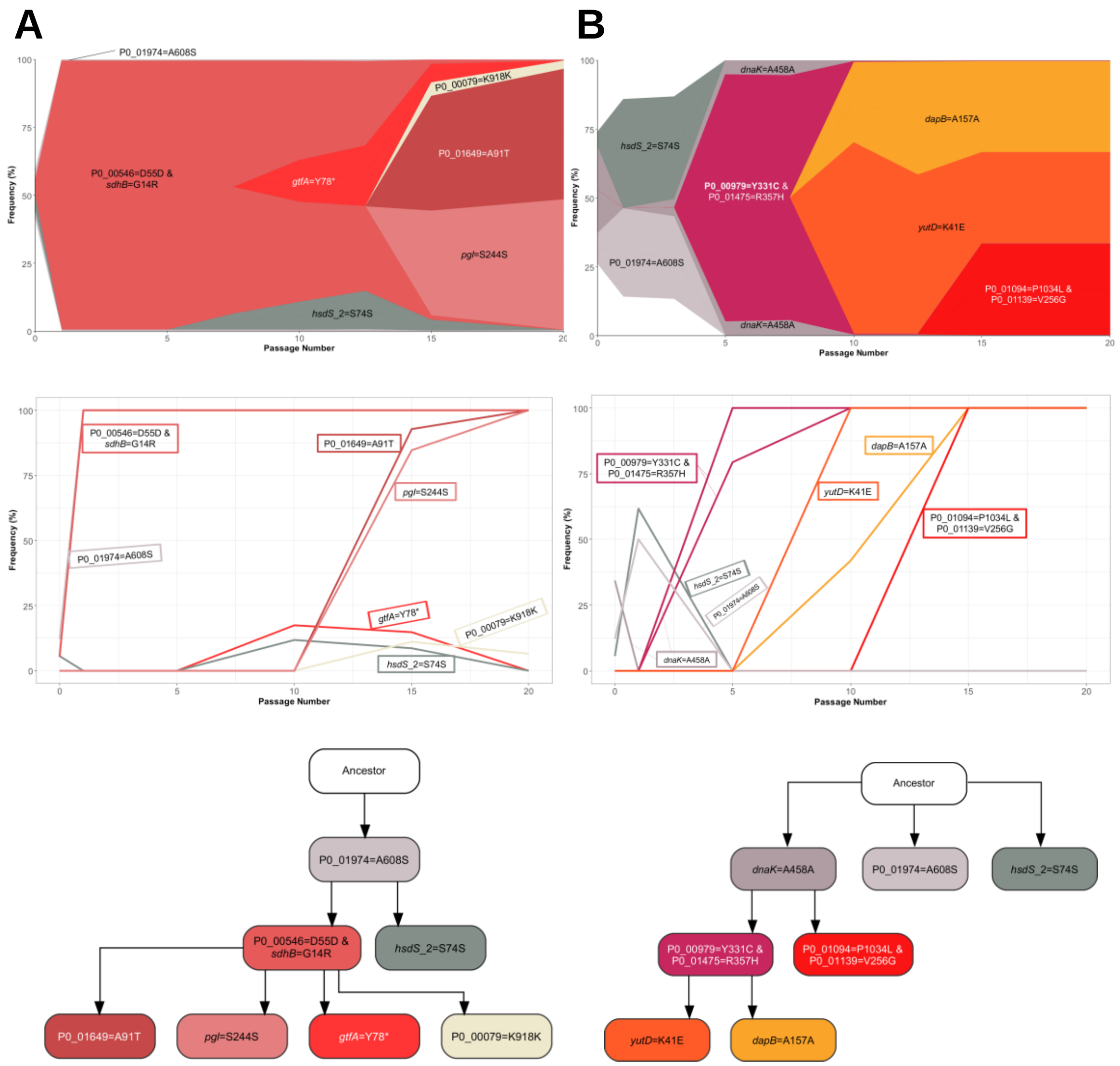


**Supplementary Figure 8. Genotype emergence and dynamics in lung passaged pneumococcal populations. (A)** Lung-passaged lineage 9. **(B)** Lung-passaged lineage 10. Panels, from top-to-bottom, are Muller plots, genotype frequency plots and genotype ancestry trees.

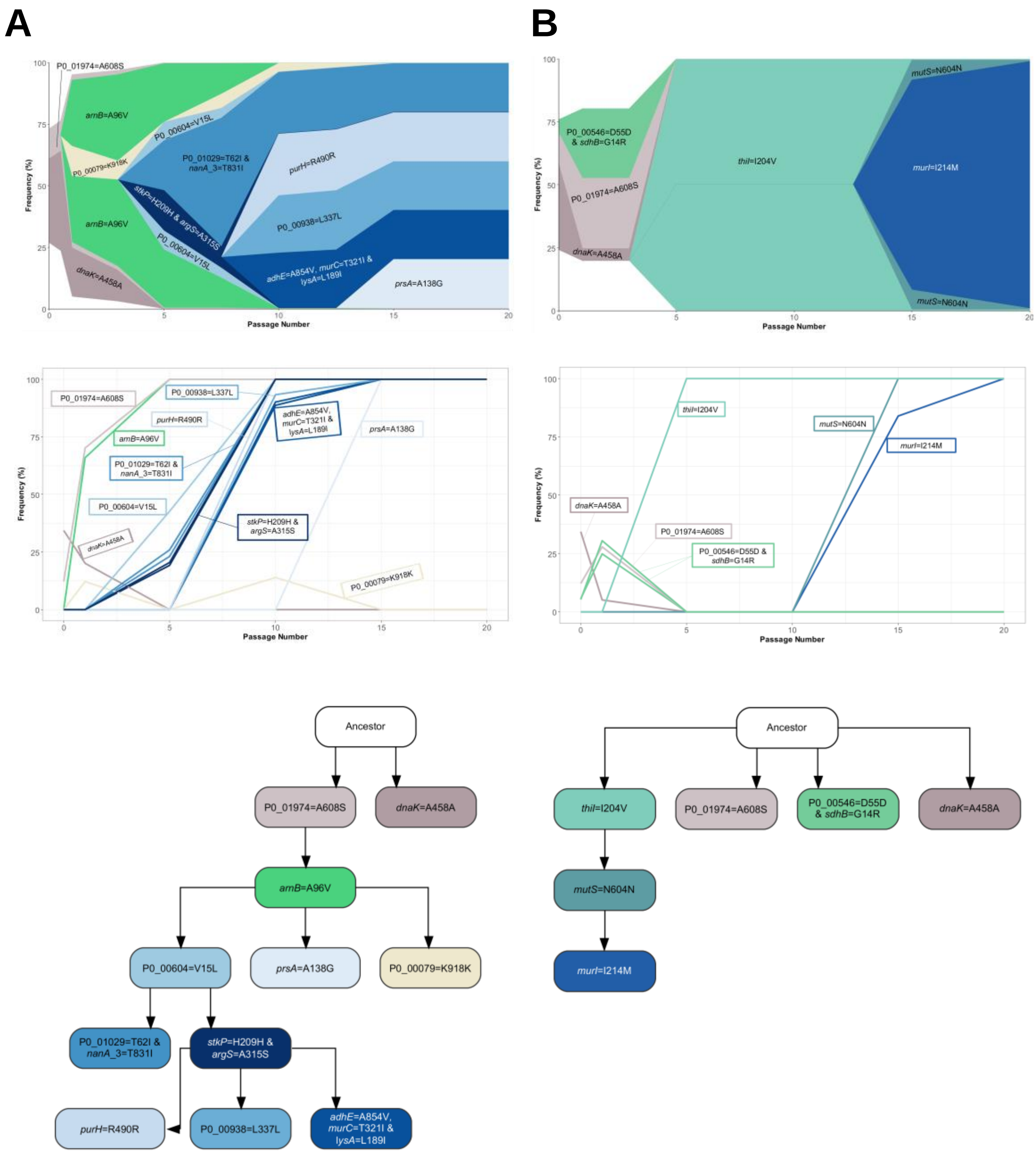


**Supplementary Figure 9. Genotype emergence and dynamics in nasopharynx passaged pneumococcal populations. (A)** Nasopharynx-passaged lineage 1. **(B)** Nasopharynx-passaged lineage 2. Panels, from top-to-bottom, are Muller plots, genotype frequency plots and genotype ancestry trees.

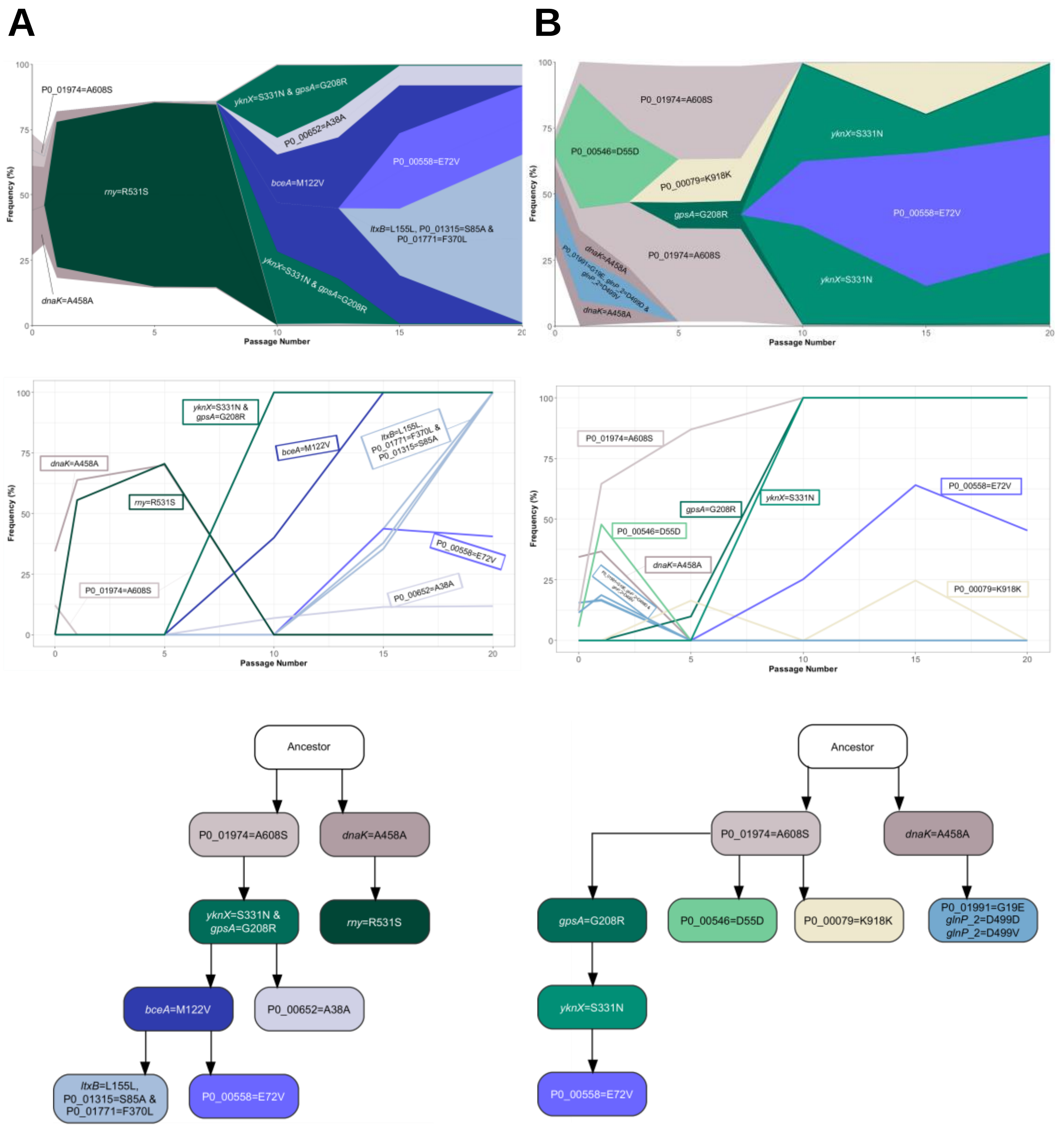


**Supplementary Figure 10. Genotype emergence and dynamics in nasopharynx passaged pneumococcal populations. (A)** Nasopharynx-passaged lineage 3. **(B)** Nasopharynx-passaged lineage 4. Panels, from top-to-bottom, are Muller plots, genotype frequency plots and genotype ancestry trees.

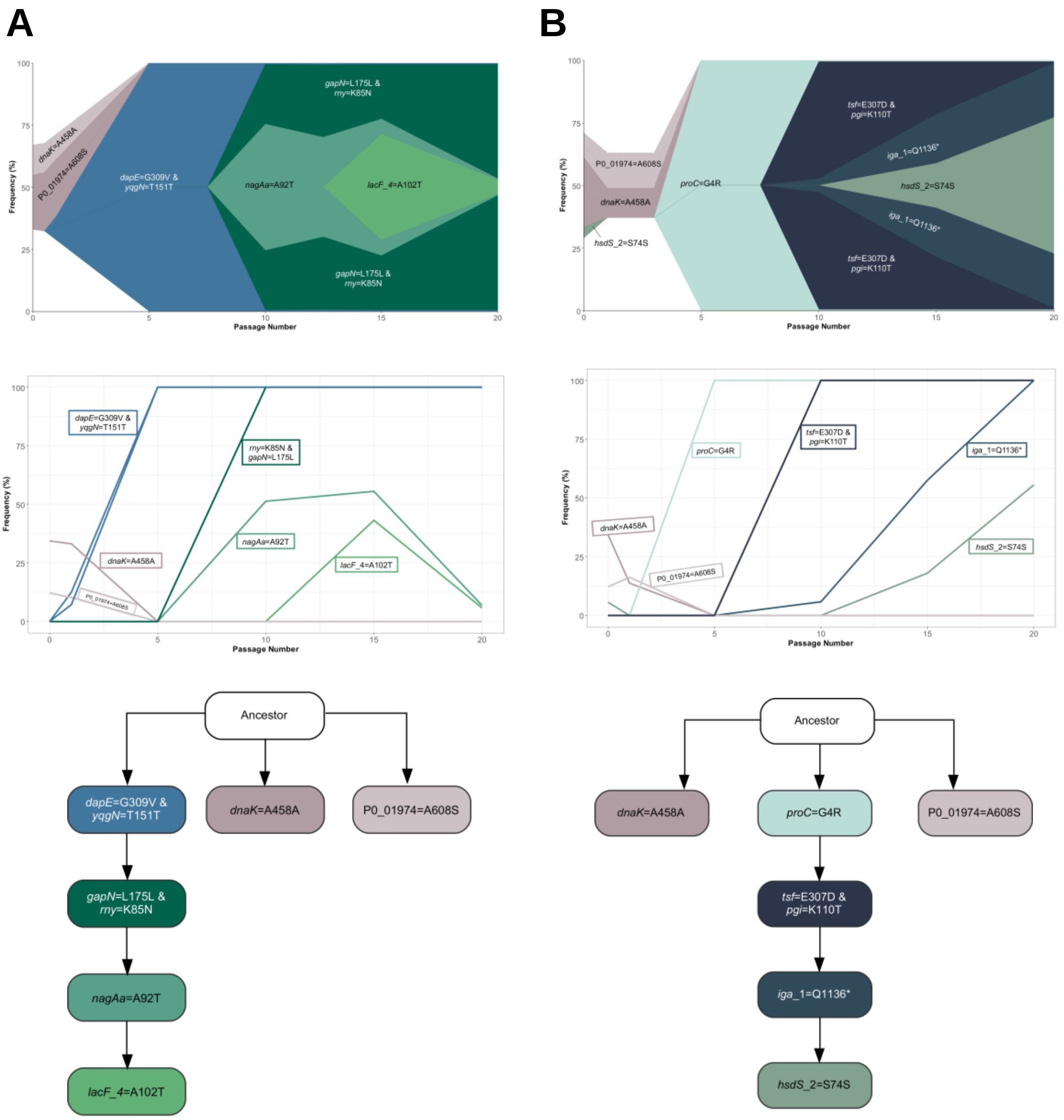


**Supplementary Figure 11. Genotype emergence and dynamics in nasopharynx passaged pneumococcal populations. (A)** Nasopharynx-passaged lineage 5. **(B)** Nasopharynx-passaged lineage 6. Panels, from top-to-bottom, are Muller plots, genotype frequency plots and genotype ancestry trees.

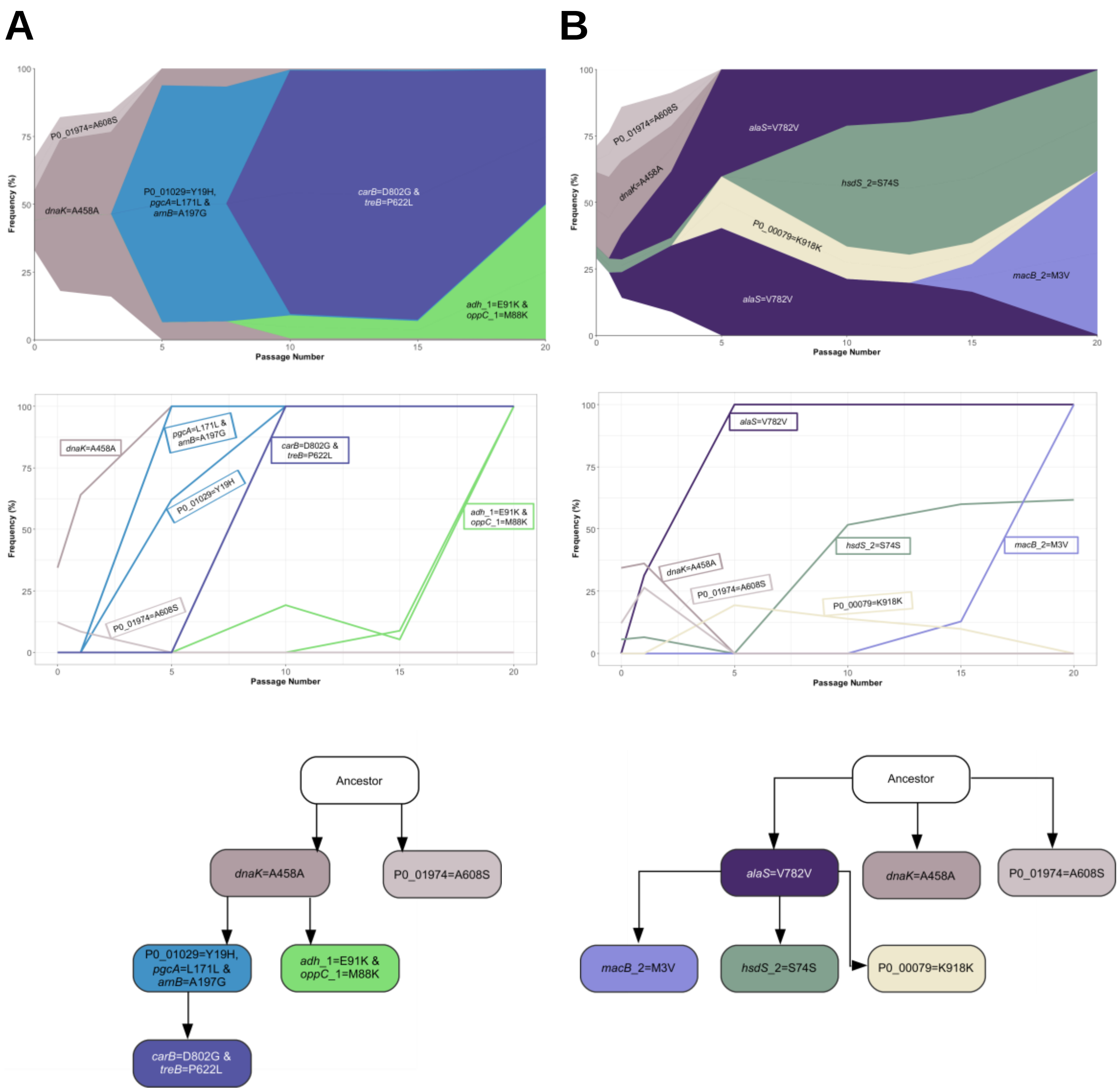


**Supplementary Figure 12. Genotype emergence and dynamics in nasopharynx passaged pneumococcal populations. (A)** Nasopharynx-passaged lineage 7. **(B)** Nasopharynx-passaged lineage 8. Panels, from top-to-bottom, are Muller plots, genotype frequency plots and genotype ancestry trees.

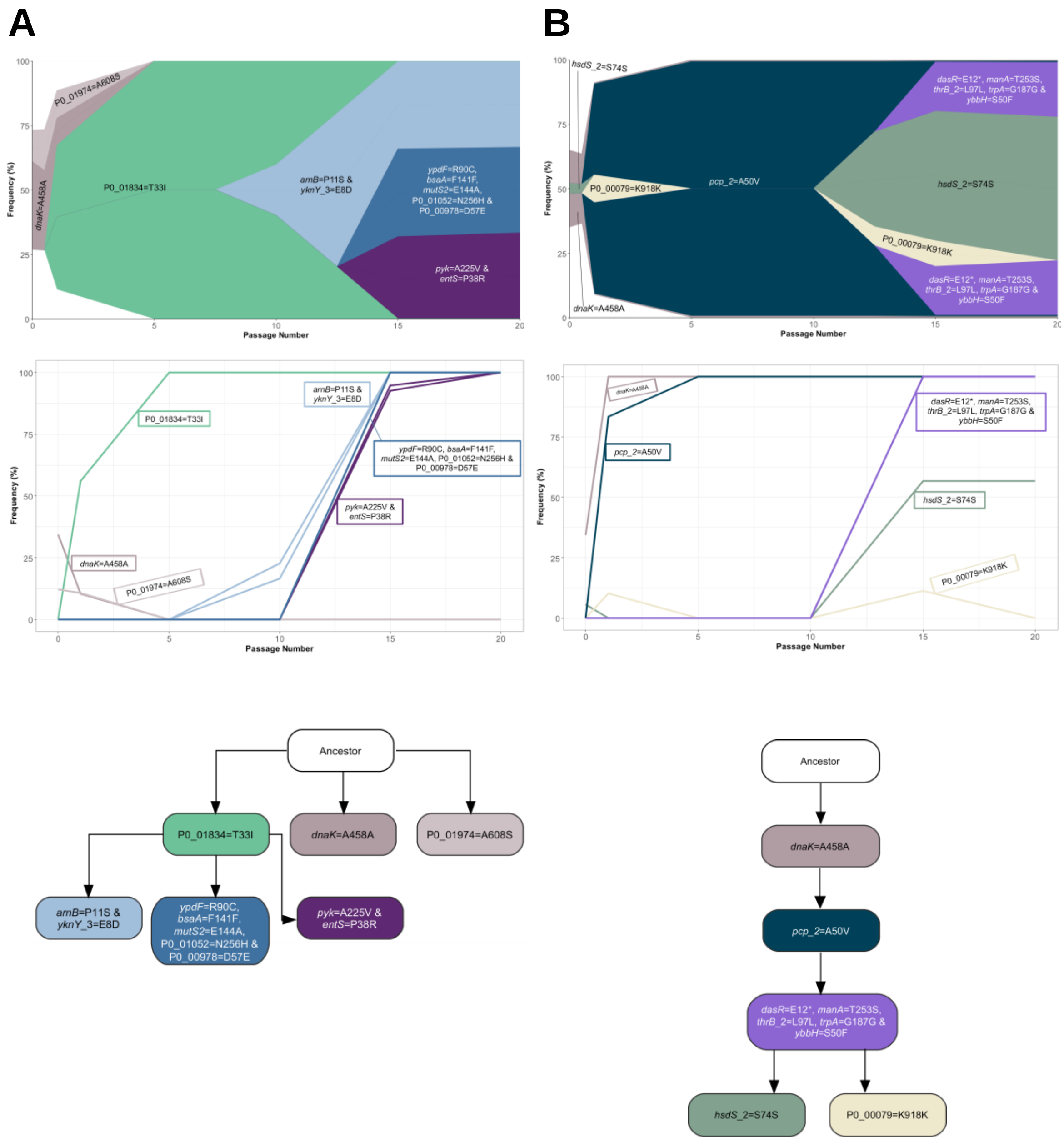


**Supplementary Figure 13. Genotype emergence and dynamics in nasopharynx passaged pneumococcal populations. (A)** Nasopharynx-passaged lineage 9. **(B)** Nasopharynx-passaged lineage 10. Panels, from top-to-bottom, are Muller plots, genotype frequency plots and genotype ancestry trees.

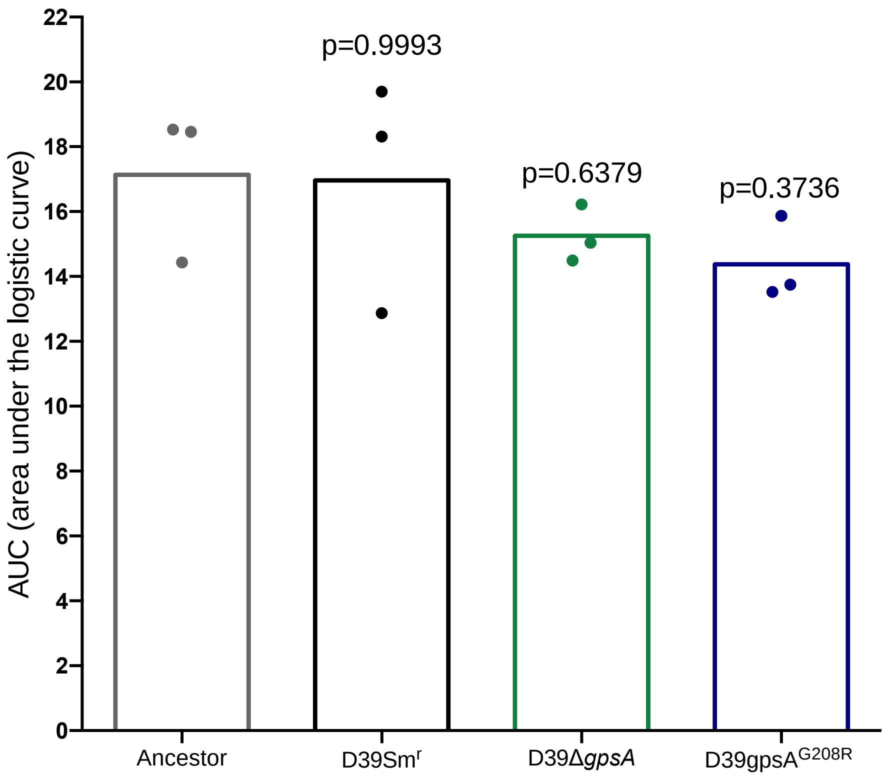


**Supplementary Figure 14. Area under the logistic curve (AUC) values of mutants on the D39 background.** The ancestor D39, the streptomycin resistant D39 control (D39Sm$_r$), the *gpsA* deletion strain (D39Δ*gpsA*) and the SNP mutant D39gpsA$_{G208R}$ were grown for 24 hours in BHI broth and AUC values were determined from the resulting growth curves. Data points are individual biological replicates, each of which is the mean of three technical replicates. P-values are from one-way ANOVA analysis with Dunnett's post-test.

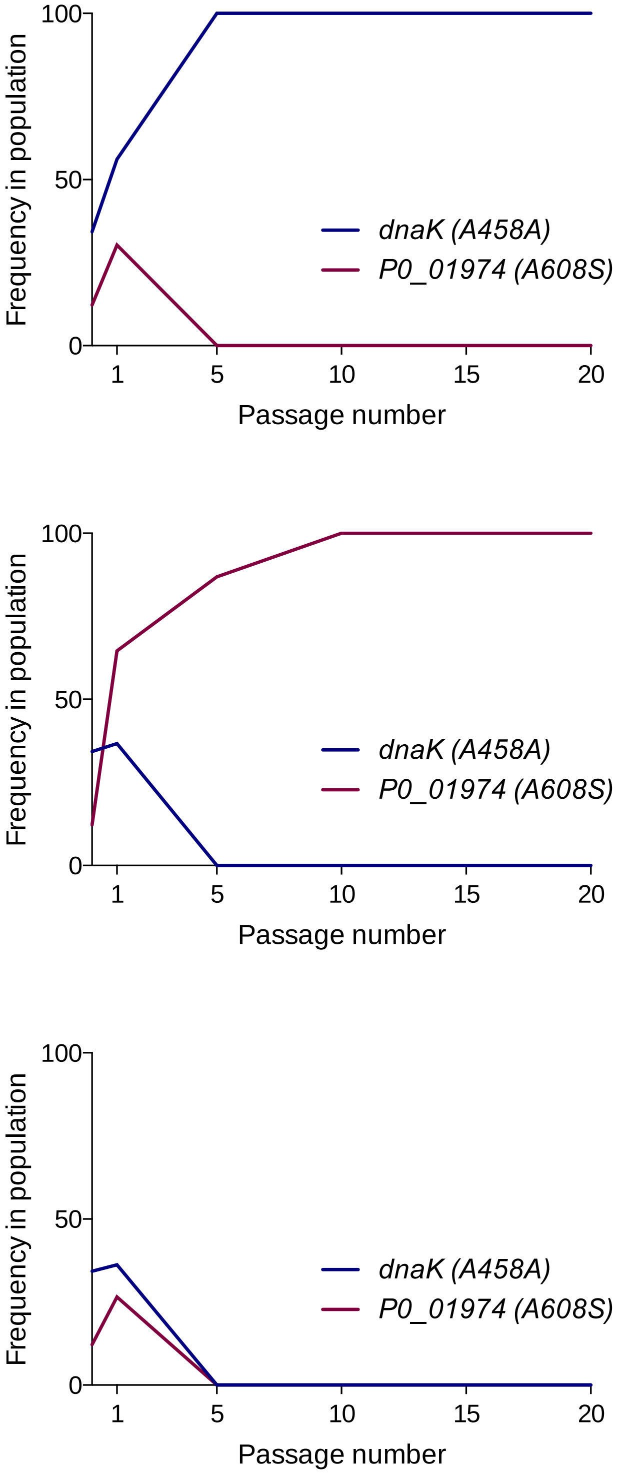


**Supplementary Figure 15. Dynamics of mutations found in the ancestor population during experimental passage.** Frequencies of synonymous *dnaK* mutation A458A and non-synonymous *P0_01974* mutation A608S over time during passage through nasopharynx and lungs. **(A)** Lung lineage 4, an example of a lineage where genotypes derived from the *dnaK* mutation-bearing portion of the ancestor drove the *P0_01974* mutation to extinction. **(B)** Nasopharynx lineage 4, an example of a lineage where *P0_01974* mutant-derived genotypes were successful at the expense of those derived from the *dnaK* mutant sub-population. **(C)** Nasopharynx lineage 8, an example of where both sub-populations were eradicated, suggesting another genotype within the ancestor population was successful.

**Supplementary Table 1. Comparison of SNPs found in mouse nasopharynx-passaged lineages and human carriage in infants.** Data from this study were compared to those from the recent publication of Chaguza *et al.* [25]. Genes shown are those with fixed SNPs identified in twenty times nasopharynx passaged populations from mice and which were also found to be targets for mutation in human carriage. SNP data from human carriage show frequencies of mutation per gene per episode of colonisation and have been normalised for gene length (per million base pairs). Episodes of colonisation lasted ~4.44 weeks (mean 7.30, range 1-48) [25].

| Gene | Present in control passage? | Number of SNPs/episode/million base pairs in human carriage | Gene product |
|---|---|---|---|
| *gpsA* | - | 216.9537286 | Glycerol-3-phosphate dehydrogenase [NAD(P)+] |
| *P0_01974* | + | 123.8729194 | hypothetical protein |
| *P0_01052* | - | 63.58701412 | hypothetical protein |
| *comEC* | - | 51.99126547 | ComE operon protein 3 |
| *thiI* | - | 50.45205037 | putative tRNA sulfurtransferase |
| *P0_00558* | - | 48.30684508 | hypothetical protein |
| *rny* | - | 41.78616878 | Ribonuclease Y |
| *nanA_3* | - | 20.02274584 | Sialidase A |
| *iga_1* | - | 18.51819576 | Immunoglobulin A1 protease |
| *bceA* | - | 16.14465612 | Bacitracin export ATP-binding protein BceA |
| *adh_1* | - | 11.8093731 | Alcohol dehydrogenase |
| *yknX* | - | 10.21643518 | Putative efflux system component YknX |
| *arnB* | + | 9.991806718 | UDP-4-amino-4-deoxy-L-arabinose--oxoglutarate aminotransferase |
| *pacS* | - | 8.198177272 | putative copper-transporting ATPase PacS |
| *pflA* | - | 7.707247896 | Pyruvate formate-lyase-activating enzyme |
| *manA* | - | 6.48516842 | Mannose-6-phosphate isomerase ManA |
| *tsf* | - | 5.887686492 | Elongation factor Ts |
| *dapE* | - | 4.461795873 | Succinyl-diaminopimelate desuccinylase |
| *pyk* | - | 4.070981636 | Pyruvate kinase |
| *carB* | - | 1.930449756 | Carbamoyl-phosphate synthase large chain |
